# Transcription machinery clustering by Integrator synchronizes histone gene expression

**DOI:** 10.1101/2023.10.07.561364

**Authors:** Feiyue Lu, Brandon J. Park, Rina Fujiwara, Jeremy E. Wilusz, David S. Gilmour, Ruth Lehmann, Timothée Lionnet

## Abstract

Numerous components of the transcription machinery, including RNA polymerase II (Pol II), accumulate in high-concentration clusters at promoters. Whether these clusters assemble on demand during transcription or constitute regulatory units remains unclear. Examining the *Drosophila* histone locus—where histone genes are transcribed exclusively during S phase—we unexpectedly found large clusters containing non-chromatin-bound Pol II and elongation factors outside S phase. When transcription is activated during S phase, P-TEFb drives accelerated turnover of cluster components. Dispersion of clusters through depletion of the Integrator complex endonuclease module causes histone transcription to have a diminished S-phase peak and instead occur ectopically throughout the cell cycle. We propose that clusters act as gatekeeping hubs by maintaining a poised machinery pool, thereby restricting gene activation to defined temporal windows.

## Introduction

Numerous regulators of eukaryotic transcription, including RNA polymerase II (Pol II) (*1–9*), co-activators (*3*, *7*, *10*), and transcription factors (*7*, *11–15*), can assemble into localized, membrane-less clusters. From early models proposing that transcription occurs at discrete, concentrated "factories", a central idea has been that these hubs play a positive role in transcription by increasing the local concentration of necessary components (*16*, *17*). Recent studies have refined this view, revealing that many of these clusters are, in fact, highly dynamic and short-lived, whose formation is often driven by the collective effect of many weak interactions between flexible protein regions known as intrinsically disordered regions (IDRs) (*3*, *4*, *10*, *11*, *18*, *19*). This dynamic model is best illustrated by the study of Pol II, whose clustering is mediated by the disordered C-terminal domain (CTD) of its largest subunit (*4*, *20–23*). *In vitro*, the Pol II CTD alone can partition into hydrogels containing other transcription factor IDRs (*20*), is sufficient to form phase-separated clusters (*21*), and can incorporate into co-activator condensates (*4*). *In vivo*, Pol II can form numerous clusters per nucleus that are liquid-like (*3*), CTD-length dependent (*21*), and correlate with transcription bursts (*2*, *5*). While this body of evidence suggests that the CTD plays a central role in organizing Pol II into hubs (*3*, *20–22*), the composition and functional interpretation of these clusters remain debated (*24*, *25*).

Directly testing the functional role of endogenous clustering *in vivo* has remained challenging due to two main limitations. First, precisely monitoring the timing of cluster assembly relative to transcriptional bursts is technically demanding. Even with state-of-the-art techniques like super-resolution and single-molecule imaging (*1–3*, *5*, *26*), the typically small size and short lifetime of endogenous clusters can obscure definitive timing due to limitations in spatiotemporal resolution, labeling sensitivity, and photostability (*27*). Second, the methods used to perturb clusters have significant caveats. Approaches that artificially induce clustering (*28–30*) often produce non-physiological effects (*31–33*). By contrast, perturbations of endogenous clusters, typically achieved by deleting IDRs of the very factors being studied, invariably reduce transcriptional output (*10*, *18*, *31–33*). Because these disordered domains often perform essential roles in recruitment and enzymatic regulation (*34–36*), their deletion might not just disrupt clustering but also cripple essential components of the transcription machinery. This potential caveat might bias towards current models in which clustering constitutes strictly an activating feature, leaving alternative regulatory functions largely unexplored despite evidence of complex, dosage-dependent effects in artificial assays (*31*, *33*).

Here, we overcome these limitations by leveraging the *Drosophila* ovarian nurse cell to investigate the regulatory function of endogenous transcription machinery clustering at the histone locus. Nurse cells are polyploid, endocycling cells with amplified genomes and large nuclear volumes (*37*), enabling excellent imaging resolution. Within these nuclei, replication-dependent histone genes - organized as a tandem array of approximately 100 copies - form the substrate of a membraneless nuclear organelle called Histone Locus Body (HLB), which concentrates the machinery required for the transcription and 3′ end processing of histone mRNAs. HLB formation requires the scaffolding protein Mxc, which assembles at histone genes and recruits various RNA and protein components to the vicinity of the locus. The transcription machinery is thought to sit at the center of the body (*38*, *39*), although whether all of the machinery is engaged in transcription within HLBs or more broadly at any loci remains to be investigated (*17*). The repetitiveness of the fly histone genes, combined with the natural polyploidy of nurse cells, creates a built-in signal amplification system. These features allow us to visualize large, microscopically distinct transcription clusters without the need for advanced microscopy. Critically, the innate coupling of histone gene activation to the S-phase transition provides a natural physiological trigger, allowing us to investigate how these clusters respond to a synchronous onset of transcription without artificial stimuli. The clarity and genetic tractability of this system also enabled us to identify a specific genetic perturbation separate from IDR mutations that disrupts the clustering of transcription components without dismantling the HLB scaffold itself. Using this approach, we find that these clusters are not simple activation sites formed by stochastic assembly of factors but instead assemble hierarchically and sequester a significant, non-productive, chromatin-unbound pool of transcription machinery. The loss of these hubs results in diminished S-phase activation and ectopic HLB transcription outside of S-phase, revealing a role for clustering in ensuring the temporal fidelity of transcription.

### Pol II, Spt5, and PAF1 form persistent clusters at HLBs before and after histone gene activation

The *Drosophila* ovarian nurse cell (NC) is a polyploid, endocycling cell that can amplify DNA to 1,000 n ploidy and reach 50 μm or more in nuclear diameter (*37*) (Fig. 1A). The most prominent Pol II clusters invariably reside on the replication-dependent histone genes (*6*, *8*, *9*, *40*), which exist in *Drosophila* as ∼100 copies at a single locus (*41*). The histone locus self-organizes into HLBs (*42*), which form one visible focus per NC in early egg chambers and multiple foci per NC in mid to late-stage egg chambers as DNA copy number increases (*43–45*) (Fig. 1B and C, marked by the HLB scaffolding protein Mxc). Thus, we leveraged the copy number repetition of the *Drosophila* histone locus and the polyploidy of NCs to enable high-fidelity imaging of transcription clusters at a native gene locus.

**Fig. 1.**
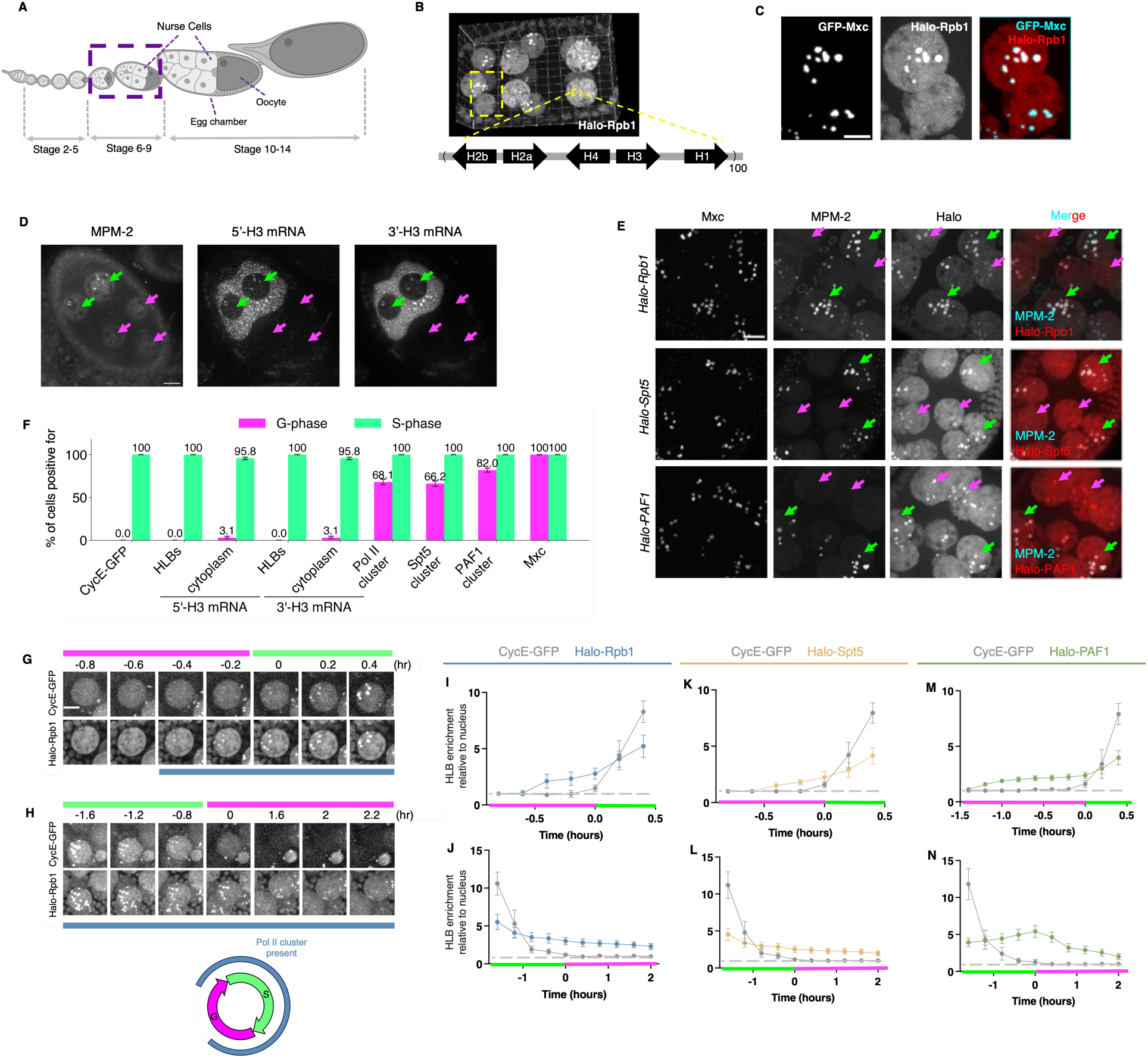
Micron-scale clusters of Pol II, Spt5, and PAF1 accumulate at histone locus bodies (HLBs) in *Drosophila* nurse cells (NCs) both during and outside of histone transcription. (**A**) Schematic of a developing ovariole. Each egg chamber contains 15 NCs and one oocyte. Boxed region indicates stages used in this study (unless otherwise noted). (**B**) 3D rendering of a stage 6 egg chamber co-expressing the HLB marker GFP-Mxc and Halo-tagged Rpb1. (**C**) Z-projection of the boxed region in (B) showing two nuclei, one of which displays distinct Pol II clusters co-localizing with HLBs. Scale bar, 5 µm. (**D**) 5′-H3 and 3′-H3 mRNA FISH signals are observed exclusively in S phase. Green and magenta arrows indicate S- and G-phase NCs, respectively; S-phase nuclei are identified by the MPM-2 enrichment at HLBs. Scale bar, 10 µm. (**E**) Clusters of Halo-Rpb1 (top), Halo-Spt5 (middle), and Halo-PAF1 (bottom) are detected in both S-phase and G-phase NCs (Halo moieties labeled with Janelia Fluor 646). Scale bar, 10 µm; Green and magenta arrows indicate S- and G-phase NCs, respectively. (**F**) Percentage of NCs positive for CycE-GFP at HLBs, 5′-H3 mRNA, 3′-H3 mRNA, Pol II clusters, Spt5 clusters, PAF1 clusters, and Mxc, stratified by endocycle phase (determined by MPM-2 staining). *n* > 300 NCs for each factor. Error bars, 95% CIs. (**G** and **H**) Representative frames from time-lapse movies of nuclei entering (G) or exiting (H) S-phase. S-phase nuclei exhibit enriched CycE-GFP at HLBs (*37*). See also fig. S2. Time 0 denotes the onset of S-phase (G) or G-phase (H). Top magenta bar marks G phase frames, top light green bar marks S phase frames; bottom color bar marks frames where Pol II is detected at the HLB. Halo-Rpb1 was labeled with Janelia Fluor 549 (*80*). Scale bar, 10 µm. (**I** to **N**) Fluorescence enrichment at HLB relative to the nucleoplasm average for Halo-Rpb1 (I and J), Halo-Spt5 (K and L), and Halo-PAF1 (M and N) at HLBs as NCs enter (I, K, M) or exit (J, L, N) S-phase. The horizontal dotted line represents the baseline nuclear level. *n* = 3 egg chambers per condition. Error bars, mean ± SEM.

Within a single egg chamber, NCs endocycle asynchronously (*37*), providing internally-matched controls for studying cell-cycle-specific transcriptional regulation. Combining MPM-2 staining to specifically mark S-phase NCs with mRNA FISH, we observed detectable cytoplasmic histone mRNAs near-exclusively in S phase, consistent with histone mRNAs being synthesized during S-phase and degraded upon exit (*45*)(Fig. 1D and F, fig. S1A to C). Histone mRNA FISH signal was also largely absent from HLBs in G phase, further demonstrating that the histone locus does not support significant levels of elongated transcripts outside of S-phase; the absence of HLB signal in G phase for a FISH probe targeting the very 5’-end of the message suggests low levels of promoter-proximally paused Pol II, although we cannot exclude the presence of Pol II paused upstream of the sequence required for probe hybridization (fig. S1A). The tight synchronization of histone gene activation with S-phase thus provides an ideal physiological context to observe how transcription clusters behave as transcription naturally begins and ends.

To test whether transcription clusters at HLBs assemble in sync with productive transcription, we analyzed Pol II together with two core transcription factors (Spt5 and PAF1), which report on distinct transcriptional stages: Spt5 associates with Pol II during promoter-proximal pausing and remains bound throughout the rest of the transcription cycle; in contrast, PAF1 is recruited later during the transition from pausing to productive elongation - the stage where active mRNA synthesis occurs - and thus serves as a marker for elongation. We generated a CRISPR-edited, homozygous viable Halo-tagged Pol II fly line expressing endogenous, chromosome-associated Rpb1 (*46*) (fig. S1D to E), and used existing CRISPR knock-in Halo-tagged Spt5 and PAF1 lines (*47*). In fixed NCs, micron-sized HLB-bound clusters of all three factors were visible in all S-phase NCs during active histone transcription, and notably, in the majority of G-phase NCs (Fig. 1E and F). Since each endocycle lasts for at least 7 hours in NCs (*37*), these transcription clusters must be present for substantial periods outside the active histone transcription window. To visualize the timing of cluster formation and decommissioning in live egg chambers, we used three fly lines, each co-expressing either of the three Halo-tagged factors with the cell cycle marker CycE-GFP (*37*), which localizes to HLBs during S-phase (*42*, *45*) (Fig. 1F). Consistent with fixed imaging results, Pol II, Spt5, and PAF1 clusters all form before entry into S-phase and persist for hours after exiting S-phase (Fig. 1G to N, fig. S2, movie S1 and S2). The lack of transcription clusters at HLBs over a fraction of G-phase is not simply due to HLB disappearance: unlike in tissue-cultured cells or early fly embryos where HLBs gradually dissolve during cell division and reform upon mitotic exit (*8*, *42*, *48*), micron-sized HLBs marked by Mxc are present in all NCs regardless of cell cycle stages (*43–45*) (Fig. 1E and F), presumably due to the lack of cell division. This suggests that the mechanisms that control transcription clusters at HLBs are separate from those that regulate HLB assembly. Collectively, these observations reveal that transcription clusters at HLBs are not merely a consequence of productive transcription, as they form prior to S-phase entry and persist well into G-phase after productive transcription has ceased.

### Clusters contain non-engaged components of the transcription machinery

To further address the nature of cluster assembly at HLBs, we asked whether promoter-proximal pausing and downstream post-pause steps (elongation and termination) are sufficient to explain clustering at HLBs. Promoter-proximal pausing is a key regulatory checkpoint in which Pol II transiently halts shortly after initiation. (*49*, *50*). At histone genes, Pol II pauses during G-phase and is released into productive elongation upon entry into S-phase (*51*). To test whether clustering depends exclusively on pause and post-pause steps, we blocked transcription initiation with triptolide (TRI) (*52*, *53*). If pausing and post-pause steps alone drive clustering, TRI should eliminate transcription clusters. CUT&RUN confirmed efficient initiation blockade, showing near-complete loss of Spt5 and PAF1 and a strong reduction in Pol II occupancy genome-wide (fig. S3A to D), including at the histone loci (Fig. 2D, F, H, and J). Because Spt5 stabilizes paused Pol II and remains associated with it through the transcription cycle, and PAF1 is recruited to Spt5-bound Pol II upon pause release and persists during elongation, their loss indicates depletion of paused and post-paused transcription species. Consistent with the loss of paused species, immunofluorescence staining showed a complete elimination of Spt5 enrichment at HLBs (Fig. 2A and G). Yet, Pol II fluorescence at HLBs decreased only partially (Fig. 2A and E), and this decrease was not due to Pol II degradation (fig. S3E and F). Since paused Pol II can remain stable for tens of minutes after initiation blockade (*52*), TRI treatment might, in principle, inhibit new Pol II accumulation while residual paused Pol II from earlier initiation events persists. To rule out this possibility, we first treated egg chambers with TRI and then monitored Pol II levels at HLBs by live imaging as cells entered S phase. This revealed the formation of *de novo* Pol II clusters at S-phase entry (although at reduced levels), supporting that clusters do not solely consist of paused Pol II but must include Pol II species insensitive to initiation inhibition (fig. S4).

**Fig. 2.**
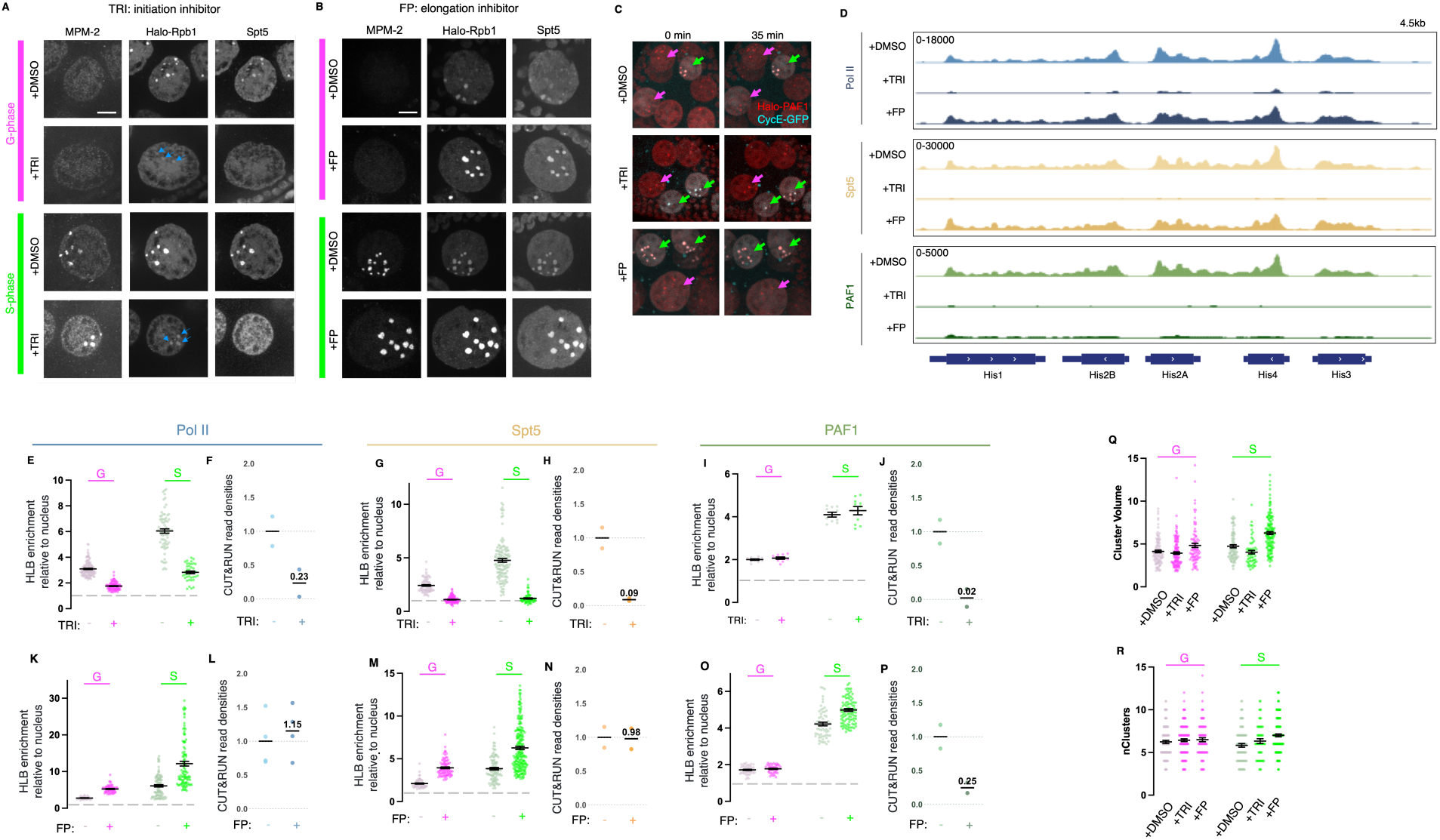
Comparison of fluorescence intensity and CUT&RUN occupancy reveals non-engaged Pol II, Spt5, and PAF1 at HLBs following transcription inhibition. (**A** and **B**) G-phase (top) and S-phase (bottom) NCs (endocycle phase demarcated by MPM-2) probed for Pol II and Spt5, treated with triptolide (TRI, A) or flavopiridol (FP, B). (**C**) Snapshots of Halo-PAF1-expressing NCs treated with DMSO (control), TRI, and FP, respectively. CycE-GFP serves as an S-phase marker. Green and magenta arrows indicate S- and G-phase NCs, respectively. Scale bars, 10 µm. (**D**) Pol II, Spt5, and PAF1 CUT&RUN occupancy at the replication-dependent histone locus in DMSO control, TRI- and FP-treated conditions. All read counts were spike-in calibrated. (**E** to **P**) Fluorescence enrichment at HLBs and average CUT&RUN occupancy over all histone gene arrays for Pol II, Spt5, and PAF1 upon transcription inhibition by triptolide (E-J) or flavopiridol (K-P). For fluorescence intensity quantifications, each dot represents one NC. The horizontal dotted line represents the baseline nuclear level. Error bars, mean ± SEM. For read density quantification, each dot represents one replicate (*n* = 4 for L; *n* = 2 for F, H, J, N, and P). (**Q** and **R**) Cluster volume (Q) and number of clusters per NC (R) in the indicated treatment conditions. Each dot represents one NC. Error bars, mean ± SEM.

We also examined PAF1, whose chromatin-bound pool was similarly depleted by TRI as measured by CUT&RUN. Strikingly, PAF1 fluorescence enrichment at HLBs was largely unchanged despite near-complete loss of CUT&RUN signal at histone genes (Fig. 2C and I). This result directly demonstrates that most PAF1 at HLBs exists as a largely chromatin-unbound or only transiently chromatin-associated (hereafter “non-engaged”) species.

To further test this conclusion, we inhibited pause release using flavopiridol (FP), an independent perturbation that blocks P-TEFb–dependent elongation. FP reduced Pol II Ser2 phosphorylation (fig. S3G to I) and strongly decreased chromatin-bound PAF1 genome-wide (fig. S3C), including at histone loci (Fig. 2D and P), consistent with effective inhibition of pause release (*54*). As exemplified at the *hsp83* locus, this blockade resulted in a depletion of Pol II, Spt5, and PAF1 signals in the gene body, while Pol II and Spt5 remained enriched near the TSS (fig. S3D). This localized accumulation aligned with the broader increase in chromatin-bound Pol II and Spt5 observed at many promoters genome-wide, consistent with previous descriptions of inhibited pause release (*52*, *53*). Yet PAF1 fluorescence at HLBs again remained largely unchanged (Fig. 2C and O), supporting the conclusions that most PAF1 at HLBs is non-engaged. FP also provided further evidence for a non-engaged Pol II pool: while Pol II and Spt5 fluorescence at HLBs increased upon FP treatment (Fig. 2B, K, and M), CUT&RUN signal at histone promoters remained largely unchanged (Fig. 2D, L, and N). Consequently, the FP-induced increase in HLB fluorescence cannot be attributed to a build-up of stably paused Pol II at the histone promoters, but instead reflects the accumulation of a non-engaged pool of Pol II and Spt5 in the vicinity of the locus. Furthermore, the sizes and numbers of HLBs were largely unaffected by blocking transcription (Fig. 2Q and R), suggesting that changes in the levels of clustered components are not simply due to changes in the HLB itself.

Together, these results demonstrate that the promoter-proximal pausing and subsequent steps of the transcription cycle are insufficient to explain Pol II clustering at HLBs. Most strikingly, nearly all PAF1 at HLBs exists as a non-engaged species, as shown by two independent perturbations. Convergent evidence from TRI, FP, and live imaging further supports the existence of a non-engaged Pol II pool.

### Pol II and Spt5 exhibit faster recovery in S phase compared to G phase, and this transition depends on P-TEFb activity

Given that transcription clusters persist at HLBs independent of productive histone transcription, we hypothesized that entry into S phase would trigger a shift in the exchange kinetics of clustered factors, reflecting a transition from a poised, sequestered state to a productive, mobile one. To test this, we performed fluorescence recovery after photobleaching (FRAP) on a single HLB in fly lines co-expressing CycE–GFP (to stage cells in G vs S phase) and Halo-tagged Pol II, Spt5, or PAF1 (Fig. 3A). Indeed, Pol II and Spt5 recovered significantly faster in S phase than in G phase at HLBs (Fig. 3B and C) but not elsewhere in the nucleus (fig. S5A and B), indicating increased exchange at HLBs upon entry into productive transcription. This difference is unlikely to reflect a general change in the HLB environment that would accelerate molecular exchange in S phase, because PAF1 recovery was slightly faster in G phase than in S phase (fig. S5C); furthermore, TRI treatment, which abolishes bound Spt5 at HLBs, results in similarly fast Spt5 recovery kinetics at HLBs and outside of HLBs (fig. S5D), suggesting that the slower recovery typically observed at HLBs is driven by specific molecular interactions.

**Fig. 3.**
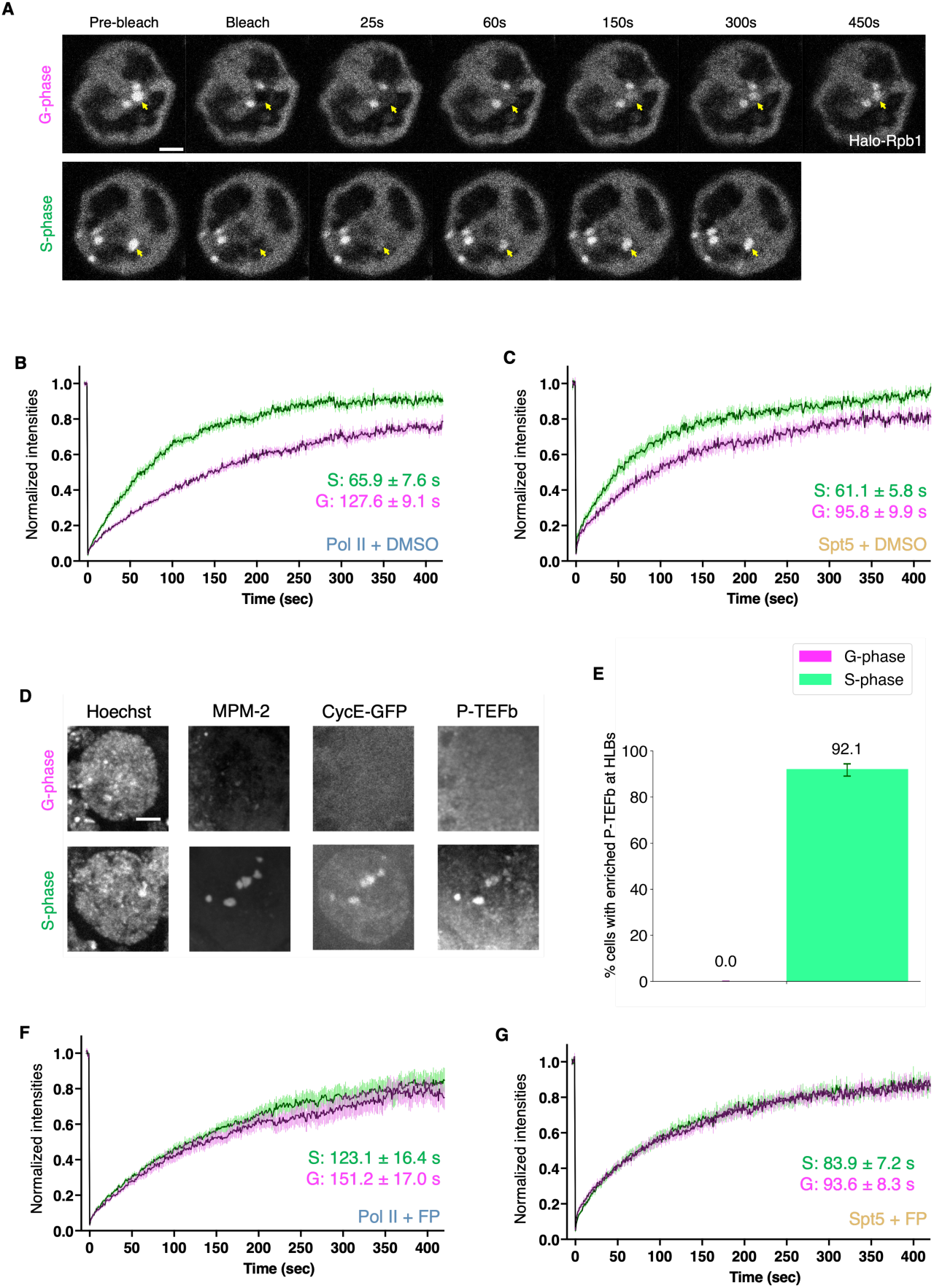
P-TEFb activity is required for the enhanced mobility of Pol II and Spt5 at HLBs during S-phase. (**A**) Example frames from FRAP experiments targeting HLB-bound Pol II clusters. Yellow arrows indicate bleached clusters. The upper G-phase cluster showed only partial recovery after 450 seconds, whereas the lower S-phase cluster showed near-complete recovery after 150 seconds. Halo-Rpb1 was labeled with Janelia Fluor 552 (JF552)(*81*). Scale bar, 5 µm. (**B** and **C**) FRAP curves of G-phase (magenta) and S-phase (green) HLB-bound Pol II (B) and Spt5 (C) clusters at HLBs in DMSO control treatment. Black lines: average curves from 7-10 nuclei across at least 5 ovaries. Vertical colored lines, SEM. A single exponential recovery curve was fitted to each factor to estimate half-lives. (**D**) Representative images of G-phase and S-phase NCs (endocycle phase demarcated by MPM-2) that express CycE-GFP and are probed for P-TEFb. Scale bar, 10 µm. (**E**) Percentage of NCs positive for P-TEFb at HLBs (S-phase vs. G-phase, as determined by MPM-2 staining). *n* = 153, 151 for G-phase and S-phase NCs, respectively. (**F** and **G**) FRAP curves of G-phase (magenta) and S-phase (green) HLB-bound Pol II (F) and Spt5 (G) clusters in FP treatment.

Because P-TEFb is highly enriched at HLBs in S phase (Fig. 3D and E), we next tested whether P-TEFb activity drives the S-phase acceleration in Pol II and Spt5 dynamics. Inhibiting P-TEFb with FP shifted the S-phase recovery kinetics of Pol II and Spt5 to resemble those observed in G phase (Fig. 3F and G, fig. S5E and F), indicating that the faster S-phase exchange at HLBs requires P-TEFb activity. To test this quantitatively, we fit the FRAP curves with a multiexponential model and found that at least three kinetically distinct populations were required (<10 s fast, 10–100 s intermediate, and >100 s slow) (fig. S6A). In G-phase, the dominant population was slow, whereas in the S-phase, the dominant population was intermediate. Upon FP treatment, S-phase kinetics became G-phase-like, with both conditions dominated by the slow population (fig. S6B). While P-TEFb is currently viewed as a regulator of the pause-release checkpoint (*55*) and a driver of hub formation (*11*, *56*), our results demonstrate that it also accelerates the turnover and exchange of the cluster constituents.

### Pol II clustering is independent of CTD length, and clustered transcription components are assembled hierarchically

Defining the functional role of clusters at HLBs requires a specific method for their disruption. We first investigated whether the Pol II CTD serves as the primary determinant of assembly, reasoning that if clustering is CTD-driven, Pol II variants with longer CTDs should exhibit increased enrichment at HLBs (*21*).To directly compare the clustering propensity of Pol II molecules containing CTDs varying in length and sequence composition, we mated an endogenously HA-tagged wild-type Rpb1 (HA-Rpb1^WT^) fly line to various FLAG-tagged Rpb1 CTD variants (*57*) such that the resulting female progeny carried one copy of each Pol II form. Contrary to the CTD-driven model, the stoichiometric ratio of FLAG-tagged CTD variants to HA-Rpb1^WT^ at HLBs did not increase with CTD length (Fig. 4A and B, fig. S7).

**Fig. 4.**
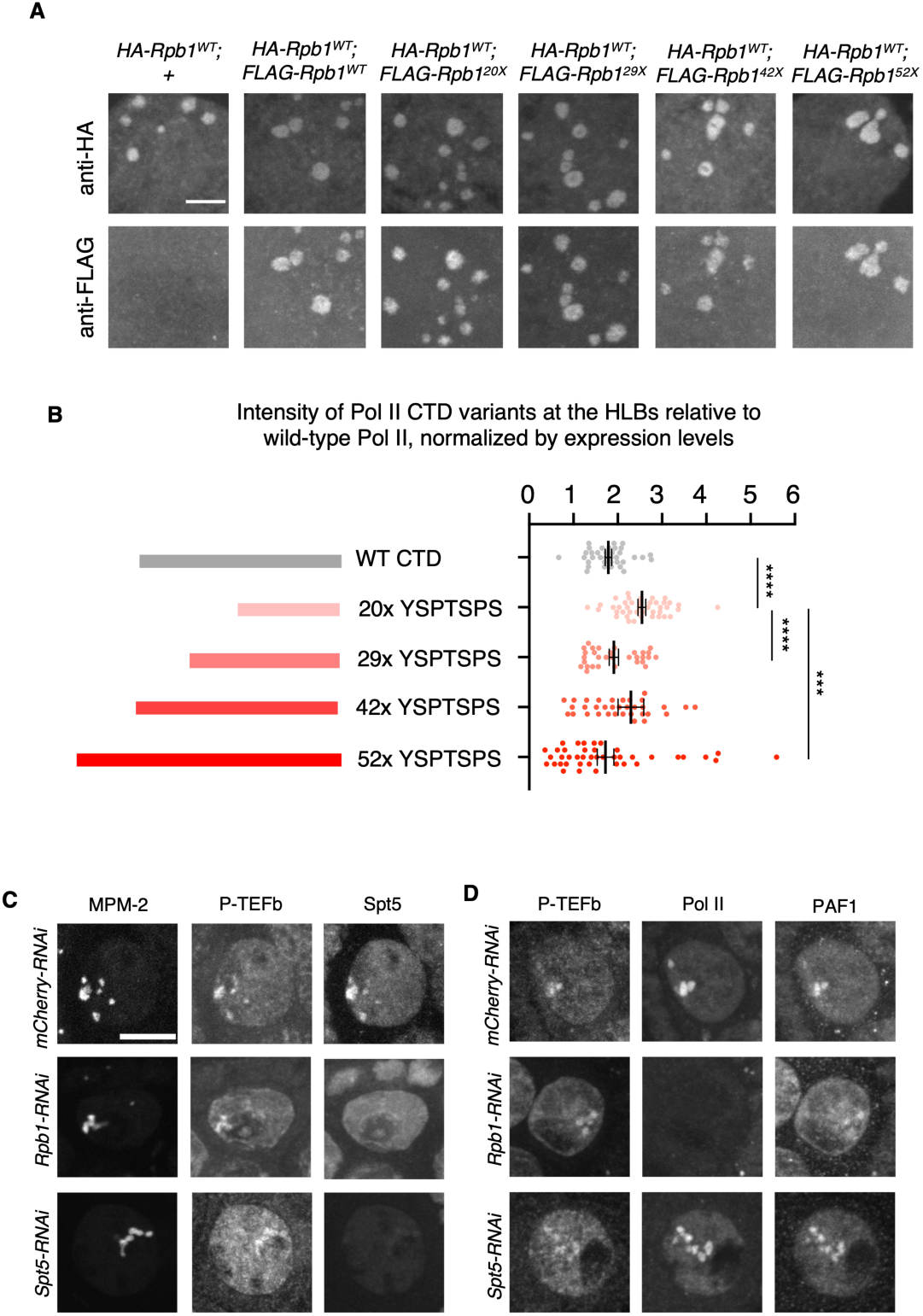
Mutational and knockdown analysis of Pol II, Spt5, and PAF1 clustering and hierarchical co-dependency. (**A**) Maximum Intensity Projections of HLB-bound Pol II clusters in the indicated genotypes. A fly line expressing an endogenously HA-tagged wild-type Rpb1 (*HA-Rpb1^WT^*) was mated to various fly lines expressing CRISPR-ed, FLAG-tagged Rpb1 CTD variants such that the resulting female progeny express one copy of each form. While the wild-type *Drosophila* CTD is composed of repeats of the YSPTSPS motif and variants of these motifs, the FLAG-tagged Rpb1 CTD variants are all consensus and consist of 20, 29, 42, and 52 repeats of the YSPTSPS motif; these are termed *FLAG-Rpb1^20x^*, *FLAG-Rpb1^29x^*, *FLAG-Rpb1^42x^*, *FLAG-Rpb1^52x^*, respectively. *FLAG-Rpb1^WT^* serves as a baseline (*57*). A mating to a fly line that does not express FLAG-tagged Rpb1 serves as a negative control. HA- and FLAG-tagged Rpb1 forms were detected using anti-HA and anti-FLAG antibodies, respectively. Scale bar, 5 µm. (**B**) Intensity of Pol II CTD variant alleles at HLBs relative to that of the wild type allele. Each dot represents the ratio of FLAG-tagged Pol II CTD variants to HA-tagged wild-type (WT) Pol II at HLBs, averaged across one nucleus (each channel normalized by nucleoplasm expression). Error bars, mean ± SEM. *n* >= 4 flies per genotype. \*\*\**P*<0.001 and \*\*\*\**P*<0.0001. See also fig. S7. (**C** and **D**) RNAi depletion of Rpb1 results in a loss of Spt5 clusters (C) but not PAF1 clusters (D). RNAi depletion of Spt5 does not eliminate either Pol II clusters or PAF1 clusters (D). Scale bar, 10 µm.

We next tested the co-dependency among cluster-associated factors. RNAi depletion of Rpb1 eliminated Spt5 clustering, but PAF1 clusters persisted. Conversely, RNAi depletion of Spt5 did not abolish Rpb1 nor PAF1 clustering (Fig. 4C and D). These outcomes are inconsistent with an independent-assembly model (in which depletion of one factor would not affect the others) or a strictly co-dependent model (in which depletion of any individual factor would collapse all clusters). Instead, they support a hierarchical assembly relationship among cluster-associated transcription regulators, whereby Pol II recruits Spt5 and PAF1 associates either upstream or independently of Pol II and Spt5. Interestingly, this hierarchy is different from the canonical stepwise recruitment to chromatin during the transcription cycle, where Pol II is recruited to chromatin first, followed by Spt5 binding, which subsequently recruits PAF1. In contrast, we show that PAF1 clustering persists upon depletion of either Rpb1 or Spt5 (Fig. 4D), and PAF1 enrichment at HLBs is detected much earlier than that of Pol II and Spt5 (Fig. 1I, K, and M). These results indicate that clustering dependencies differ from the established temporal order of chromatin recruitment.

### IntS11 is required for clustering independently of HLB scaffold integrity

To identify potential regulators of clustering, we reasoned: (i) they should function early in the transcription cycle, consistent with evidence that clusters are apparent in G phase; and (ii) they should have the potential to both activate and repress transcription, consistent with a model in which clustered factors are poised for synchronous activation in S phase. The Integrator complex meets both criteria. The Integrator is a multi-subunit complex that regulates promoter-proximal transcription and contains two functional modules: a phosphatase and an endonuclease, associated with nascent RNA cleavage and restriction of Pol II activity (*58–61*). RNAi depletion of the Integrator endonuclease module subunits IntS11 or IntS9 resulted in a complete loss of visible HLB-bound Pol II, Spt5, and PAF1 clusters (Fig. 5A to C, E to G, and fig. S8A to E, G to I). Importantly, the loss of transcription clusters upon IntS11 or IntS9 RNAi is not simply due to a global loss of HLB structure, as the scaffolding protein Mxc remained present at HLBs (Fig. 5D, H to J, fig. S8F, J to L). Nor is it due to RNAi-induced cell death, as IntS11 RNAi and IntS9 RNAi egg chambers progressed through the stages analyzed in this study (fig. S8M). RNAi depletion of IntS8, a key bridging subunit connecting the phosphatase module to the rest of the Integrator (*62*, *63*), resulted in increased phosphorylation on the Pol II CTD Ser5, consistent with the loss of phosphatase activity (*61*), yet this did not reduce Pol II levels at HLBs (fig. S9). Collectively, these results establish the Integrator endonuclease module, but not its phosphatase module, as an essential regulator for clustering at HLBs.

**Fig. 5.**
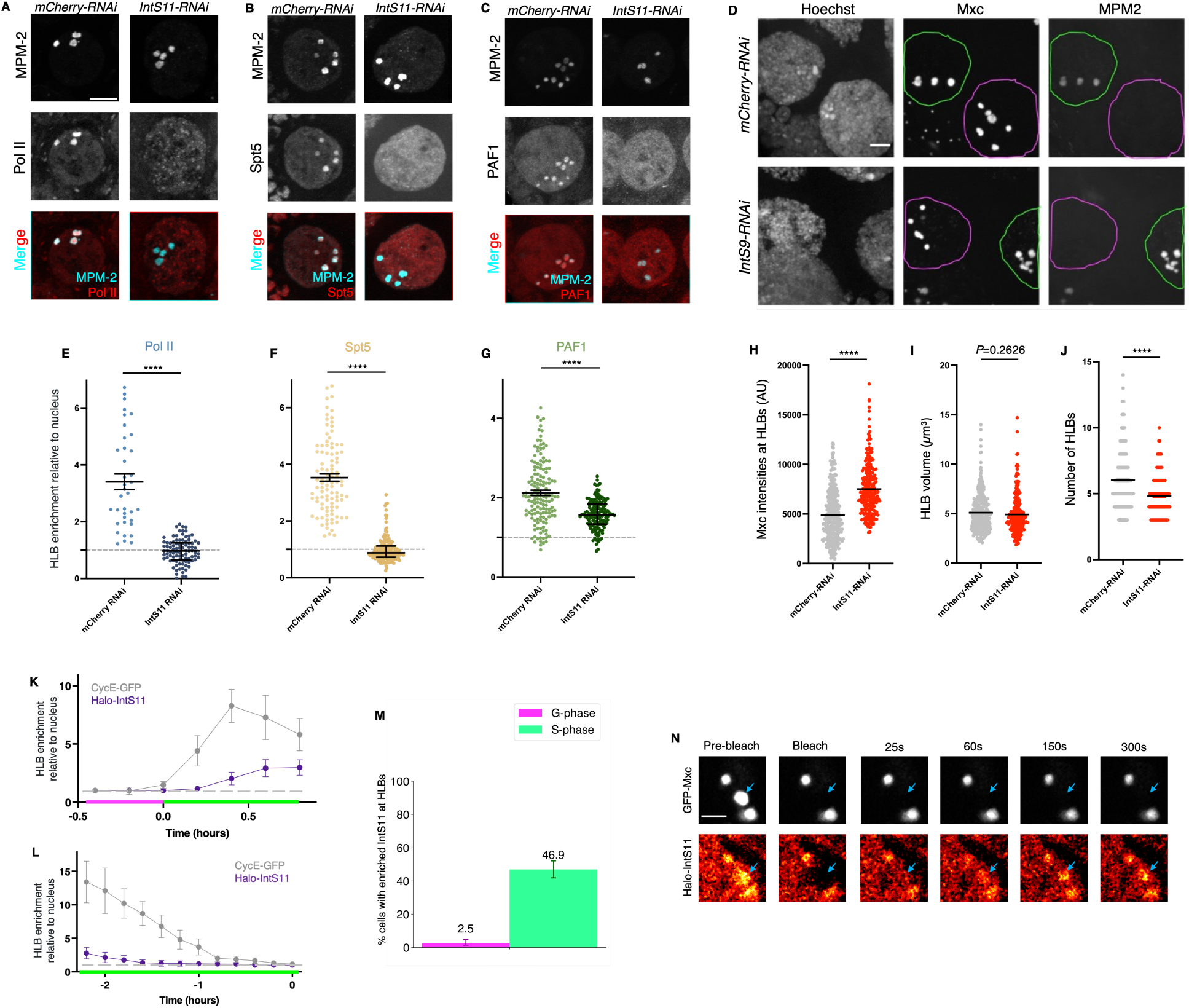
IntS11 is required for clustering at HLBs through recruitment and turnover kinetics that are distinct from those of structural scaffolds. (**A** to **D**) RNAi depletion of IntS11 results in a loss of Pol II (A), Spt5 (B), and PAF1 clusters (C) at HLBs, whereas Mxc levels are not reduced (D). mCherry-RNAi serves as a negative control. Scale bars, 10 µm. (**E** to **J**) Quantification of Pol II (E), Spt5 (F), PAF1 (G), and Mxc (H to J) at HLBs. Pol II, Spt5, and PAF1 (E to G) intensities at HLBs (demarcated by MPM-2) were normalized to nucleoplasm levels. The horizontal dotted line represents the baseline nuclear level. For Mxc (H to J), HLB intensity, HLB volume, and number of HLBs per NC were measured based on Mxc staining (D). Each dot represents one NC. Error bars, mean ± SEM. All data were stage-matched and acquired from at least 6 flies per condition. \*\*\*\**P* < 0.0001. (**K** and **L**) Fluorescence intensity of Halo-IntS11 at HLBs as NCs enter (K) or exit (L) S-phase. See also fig. S8N and O. The horizontal dotted line represents the baseline nuclear level. *n* = 3 egg chambers per condition. Error bars, mean ± SEM. (**M**) Percentage of NCs enriched for Halo-IntS11 at HLBs, stratified by endocycle phase (determined by MPM-2 staining). *n* = 200 and 192 nuclei for G-phase and S-phase, respectively. Error bars, 95% CIs. (**N**) Representative FRAP experiment in a nurse cell co-expressing GFP-Mxc and Halo-IntS11, targeting Mxc and IntS11 at HLBs. The blue arrows indicate the bleached cluster. GFP-Mxc (top) does not show any recovery, whereas Halo-IntS11(bottom) shows partial recovery. Halo-IntS11 was labeled with JF552. Scale bar, 5 µm.

We next tested whether IntS11 could act as a direct scaffold for cluster-associated factors. If IntS11 were a scaffold, it should become enriched at HLBs either before or concurrent with clustered Pol II/Spt5/PAF1, and remain present throughout the clustered state. Instead, IntS11 was not detectably enriched at HLBs until after entry into S phase and did not persist for the entirety of S phase (Fig. 5 K to M, fig. S8N and O), in contrast to other cluster-associated factors, which are present for substantial periods of G phase and persist throughout S phase (Fig. 1 F to N, fig. S2). Moreover, unlike the canonical HLB scaffold Mxc, which shows undetectable FRAP recovery within 5 min, Halo-IntS11 partially recovered after photobleaching, suggesting a dynamic association (Fig. 5N). Together, these results indicate that IntS11 is required for clustering but does not function as a canonical, stably associated HLB scaffold, and that transcription clusters can be perturbed while overall HLB scaffold integrity is preserved.

### Disrupted clustering is linked to a diminished S-phase peak and ectopic G-phase expression of histone genes

Because Integrator depletion disrupts clustering while leaving the HLB scaffold intact (Fig. 5A–J and fig. S8C to L), it provides a means to examine the transcriptional consequences of cluster loss. If clustering simply enhanced transcription, then loss of clustering upon Integrator depletion should markedly reduce histone gene expression. To test this, we visualized histone expression by RNA FISH against H3 mRNA. In control egg chambers, S-phase NCs showed robust H3 signal at HLBs and in the cytoplasm, whereas G-phase NCs were essentially signal-free, consistent with prior work (*45*) (Fig. 6A, fig. S10A). RNAi depletion of IntS11 or IntS9 resulted in a narrower range of H3 mRNA expression, with a reduced higher quartile indicative of decreased levels of induction and an increased lower quartile compared to control (Fig. 6B, fig. S10B). Because an elevated lower range could reflect altered cell-cycle composition, we restricted the analysis to G-phase NCs. Strikingly, most G-phase NCs were H3 mRNA-positive after IntS11 or IntS9 depletion, and the signal often included a nascent H3 focus at HLBs, consistent with aberrant activation of histone transcription during G phase (Fig. 6C and D, fig. S10C). Notably, P-TEFb enrichment remained S-phase-specific after IntS11 or IntS9 knockdown, ruling out aberrant P-TEFb recruitment as the cause for leaky G-phase expression (fig. S8P and Q), while EdU incorporation verified that cell-cycle staging remained highly accurate across genotypes (fig. S8R).

**Fig. 6.**
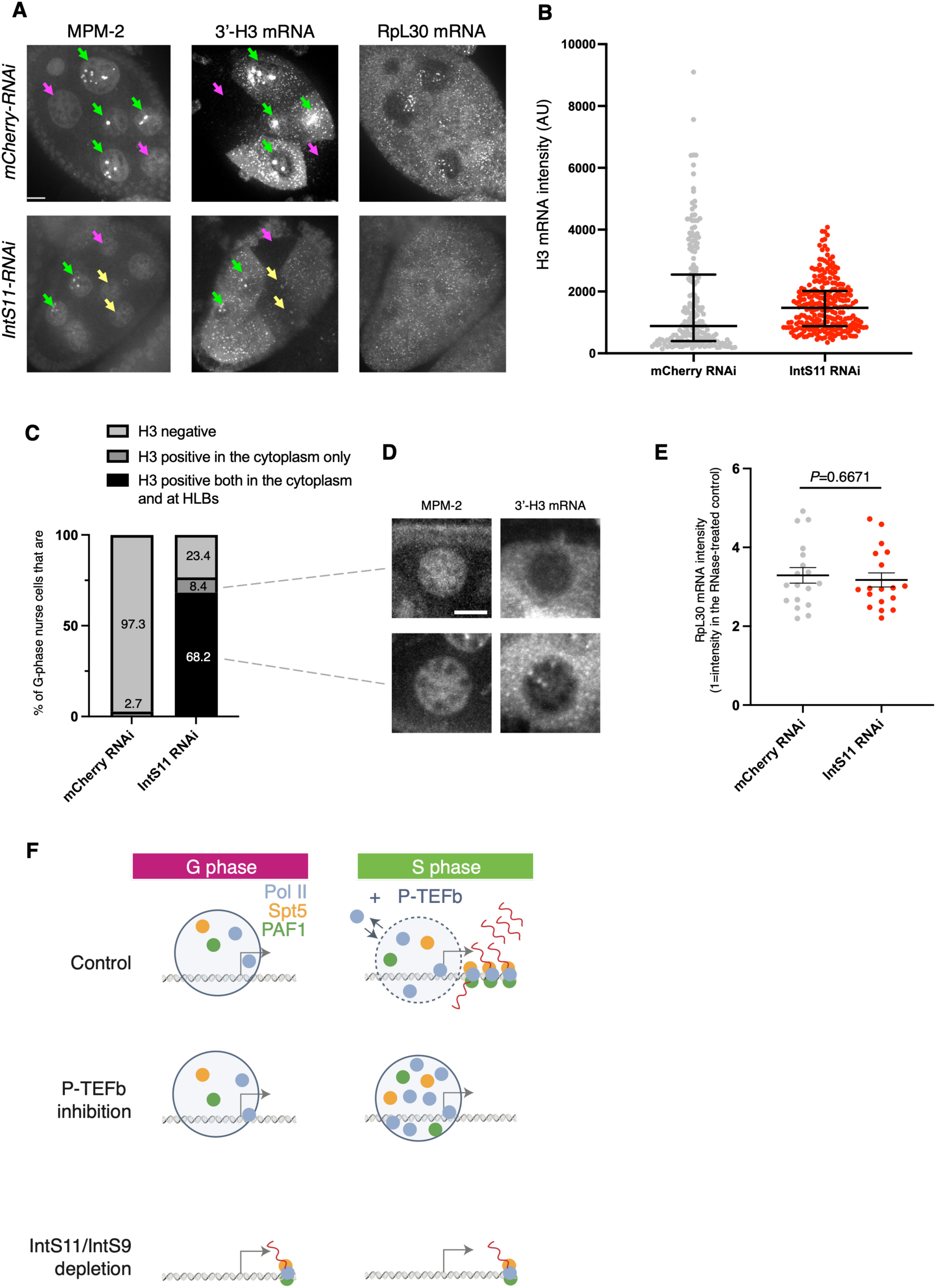
Depletion of IntS11 results in reduced induction and out-of-phase histone transcription. (**A**) Egg chambers probed with MPM-2 (S-phase marker), 3’-H3 mRNA, and RpL30 mRNA (control). mCherry-RNAi serves as a negative control. Green arrows indicate S-phase NCs, whereas magenta and yellow arrows indicate G-phase NCs lacking and containing H3 mRNA, respectively. Scale bar, 10 µm. (**B**) Cytoplasmic H3 mRNA intensity. Each dot represents one NC. Bars indicate quartiles. *P*<0.00001, estimated by bootstrapping of the 10^th^ and 90^th^ percentiles. (**C**) Percentages of MPM-2-negative NCs with no H3 mRNA signal, H3 mRNA in the cytoplasm only, and H3 mRNA both in the cytoplasm and at HLBs. *n* > 300 NCs for each genotype. (**D**) Representative images showing a G-phase NC with H3 mRNA in the cytoplasm only (top) and a G-phase NC with H3 mRNA in both the cytoplasm and at HLBs (bottom). Scale bar, 10 µm. (**E**) Cytoplasmic RpL30 mRNA FISH intensity. Each dot represents the average mRNA level in one egg chamber. Error bars, mean ± SEM. (**F**) Model of clustering-regulated transcription at replication-dependent histone genes. In normal conditions (top), transcription components (Pol II, Spt5, and PAF1) form clusters near histone promoters during G-phase but remain inactive. Pol II and Spt5 exhibit a slow turnover rate. Upon entry into S-phase, P-TEFb activity licenses these clusters, promoting the assembly of productive elongation complexes and increasing the recovery rate of Pol II and Spt5 by FRAP. P-TEFb inhibition by flavopiridol (FP) (middle) blocks the licensing required for productive transcription. Consequently, Pol II and Spt5 in S-phase maintain a slow, G-phase-like recovery rate, indicating prolonged residence time within the cluster. Loss of the Integrator endonuclease subunits (bottom) abolishes clustering. As a result, in G-phase, transcription components bypass the P-TEFb-gated licensing step and become elongation-competent immediately upon recruitment. This results in aberrant "leaky" histone transcription during G-phase and a failure to properly build up the Pol II levels required to induce high-level transcription during S-phase.

To test whether this elevated G-phase H3 mRNA signal could reflect a genome-wide increase in transcription upon Integrator loss, we first measured cytoplasmic RpL30 mRNA and observed a partial decrease upon IntS9 RNAi and no change upon IntS11 RNAi, arguing against a global disruption of mRNA levels (Fig. 6A and E, and fig. S10A and D). We next tested whether loss of HLB-associated transcription clusters impairs histone pre-mRNA 3′ end processing, an outcome that could occur if cluster loss compromises HLB functional/structural integrity. Using an RNA FISH probe downstream of the H3 mRNA cleavage site (DS-H3) to detect unprocessed transcripts (*42*, *45*) we found that the DS-H3:3′-H3 signal ratio did not increase at HLBs or in the cytoplasm upon IntS11 or IntS9 RNAi (fig. S11), arguing that histone mRNA processing is not impaired upon Integrator endonuclease depletion.

To narrow down when ectopic expression arises, we staged NCs more precisely by EdU pulse labeling followed by a 75-min chase and MPM-2 staining, enabling separation into four cell-cycle phases (fig. S12A). In controls, early S-phase NCs ranged from H3-negative to strongly H3-positive, consistent with histone transcription initiating after S-phase entry. In contrast, after IntS11 RNAi, early S-phase NCs uniformly showed detectable H3 mRNA signal (fig. S12B), indicating that histone transcription is initiated prematurely. Because H3 mRNA is already detectable at the earliest S-phase stage, this argues that the “leaky” histone transcription seen after Integrator loss is not simply due to a failure to shut down transcription during S-phase exit. Together, these results support a model in which Integrator-dependent clustering enforces cell-cycle–coupled histone transcription by restricting productive activation to S phase (Fig. 6F).

## Discussion

Clustering of the transcription machinery is thought to boost transcription by forming hubs that locally concentrate components to promote gene expression (*16*, *17*). However, our findings in the *Drosophila* nurse cell reveal a more complex role for these assemblies in enforcing the temporal fidelity of gene activity. This discovery was facilitated by the well-defined nature of replication-dependent histone transcription, where we find that large clusters of Pol II, Spt5, and PAF1 form at HLBs well before the onset of S-phase and persist long after transcription has ceased. While in some contexts clustering can be accounted for by elongating species (*25*, *64*), we observe the persistence of clusters outside the active transcription window. This suggests that clustering represents a fundamentally distinct, regulated state with a key function in ensuring the temporal fidelity of the transcription machinery release (Fig. 6). By identifying the Integrator endonuclease as a key regulator of clustering, we move beyond recent descriptions of poised species (*51*, *65*) and provide a means to perturb the gatekeeping function of the hub.

Our investigation uncovers that clusters sequester a significant pool of non-engaged machinery, suggestive of a regulatory layer physically decoupled from promoter-proximal pausing (*49*, *50*), a well-established checkpoint for regulating the timing of gene activation by stabilizing Pol II molecules already engaged with the DNA template. The persistence of Pol II and PAF1 clusters following initiation blockade demonstrates that the physical integrity of this cluster does not depend on the continuous production of paused or elongating species, allowing the maintenance of a pre-assembled reservoir independently of the canonical transcription cycle. This discrepancy mirrors recent observations that CUT&RUN selectively captures long-lived, DNA-bound species while failing to detect weak interactions that drive protein colocalization in live cells (*66*). Beyond its established role in pause release (*55*) and hub formation (*11*, *56*), P-TEFb activity regulates the kinetics of this reservoir by modulating the exchange between the clustered components and the surrounding pool (Fig. 6F, middle).

The assembly of transcription clusters at the HLB appears to follow a hierarchical logic rather than a purely stochastic assembly process. While clustering mediated by the Pol II CTD has been shown to scale with CTD length (*20*, *21*), we find that Pol II recruitment to the HLB does not. This suggests that for poised hubs, recruitment is likely governed by a specific molecular hierarchy rather than purely homotypic interactions. Indeed, we observe that the elongation factor PAF1 arrives at the HLB much earlier than Pol II or Spt5 and remains enriched even when they are depleted. Similar hierarchical assemblies have been observed for pioneer factors (*15*) and coactivators (*3*); however, our data demonstrate that PAF1, a factor traditionally associated with elongation, can also function as an early-recruited component. This finding echoes proteomic evidence identifying the PAF1 complex as a prominent constituent of coactivator hubs (*19*). By resolving this hierarchy at a native gene locus in real time *in vivo*, our data thus suggest that PAF1 may help establish the structural foundation of a transcription hub long before productive transcription begins.

We identify the Integrator endonuclease as a key regulator: its depletion dissolves transcription clusters without dismantling the underlying HLB scaffold, enabling us to test cluster function. While traditionally viewed as a promoter-proximal terminator that evicts paused Pol II (*58–61*, *67*), our data reveal a previously unappreciated requirement for the Integrator in the assembly of transcription clusters. Whether this reflects a direct role or is an indirect consequence of Integrator’s established function in promoter-proximal termination remains to be determined. While we cannot rule out that, in principle, the Integrator prevents G-phase histone expression by degrading nascent transcripts, the observation that IntS11 is not enriched at HLBs until entry into S phase argues against this scenario. Rather, our findings are consistent with the Integrator endonuclease acting as an upstream regulator of a scaffold establishing the foundational molecular environment required for the subsequent partitioning of client species (PAF1, Pol II, Spt5, and possibly others) (*68*, *69*). We speculate that the scaffold may contain non-coding RNAs, which are primary targets of Integrator-mediated processing and termination (*58*, *70*). Our observation that *IntS11* RNAi leads to the appearance of ectopic Pol II puncta away from the HLB (Fig. 5A) mirrors findings in human cells, where Integrator depletion was similarly found to enhance Pol II clustering at specific loci (*71*). We speculate that these puncta may represent Pol II complexes that fail to undergo proper termination and thus remain persistently arrested at non-coding promoters. Furthermore, because Integrator endonuclease depletion reduces transcription bursting broadly and independently of pausing (*72*), the loss of clustering at the HLB may reflect a general requirement for the complex in establishing functional transcription hubs at early transcriptional steps (*73*).

We propose that clustering functions as a molecular dam: in normal conditions, clustering sequesters Pol II during G-phase, preventing its transition into elongation and establishing a concentrated reservoir of transcription machinery. This reservoir is then rapidly released in S phase through P-TEFb activity, which serves as a molecular gate (Fig. 6F, top). Depletion of the Integrator endonuclease abolishes the dam, allowing Pol II to bypass P-TEFb-mediated licensing and become elongation-competent immediately upon recruitment. This results in aberrant, leaky G-phase expression and also in a failure to accumulate the Pol II reserves required for robust S-phase induction (Fig. 6F, bottom).

The dam analogy provides a plausible framework to reconcile the conflicting roles of the Integrator complex as both an activator (*60*, *71*, *74*, *75*) and a repressor (*76*, *77*) of transcription. A key prediction of this model is that IntS11 depletion should enable the transcription machinery to bypass P-TEFb-mediated licensing. This prediction aligns with genome-wide observations that highly expressed genes tend to decrease while lowly expressed genes increase upon the loss of IntS11 (*61*, *73*). Our findings at the histone locus mirror these bivalent effects across different cell cycle phases: in the absence of the Integrator-mediated dam, genes with normally low P-TEFb activity (where the gate is shut) experience leaky, ectopic expression, while genes normally highly active (where the gate is open) fail to accumulate the concentrated Pol II reserves required for robust induction. Ultimately, these results challenge the prevailing model that clusters form to promote transcription and instead redefine them as active regulatory hubs that enforce temporal fidelity through sequestration and licensed release. This gatekeeping framework conceptually mirrors the phenomenon of promoter-upstream Pol II ‘docking’ observed at inactive growth and developmental genes (*78*, *79*), suggesting a broader metazoan strategy in which uninitiated intermediates are leveraged to maintain strict temporal repression while remaining poised for synchronous activation. Our discovery adds a new dimension to the role of the Integrator complex and provides a conceptual framework for investigating whether similar regulators coordinate rapid, high-magnitude transcriptional responses elsewhere in development or tissue homeostasis.

## Acknowledgments

We thank John Lis and Philip Versluis (Cornell) for generating the Halo-IntS11 fly line and sharing the Halo-Spt5 and Halo-PAF1 fly lines. Robert Duronio (UNC-Chapel Hill) for the GFP-Mxc fly line and anti-Mxc antibody, Elizabeth Gavis (Princeton) for the GFP-CycE fly line, Akira Nakamura (RIKEN) for anti-CycT antibody, Stefano Di Talia (Duke) for discussion on the DS-H3 HCR FISH probe design, Luke Lavis (Janelia) for the JF dyes. Eric Wagner (URMC), Yu Cai (NUS), and Patrick O’Farrell (UCSF) for critical reagents, Yujun Chen (K-State) for *ex-vivo* ovary culture advice, Nicholas Mamrak for designing the 5’-and 3’-H3 HCR FISH probes, Xiaona Tang (JHU) for CUT&RUN suggestions, Matthew Maurano for CUT&RUN library sequencing support. Chengye Feng (Yale) for FRAP analysis advice, NYU Langone Health Microscopy Laboratory (RRID: SCR_017934) Michael Cammer and Yan Deng for consultation and assistance with optical microscopy services, the TRiP at Harvard Medical School for providing various RNAi fly stocks used in this study, the Lionnet and Lehmann lab members for comments.

## Author contributions

F.L. conceived the project. T.L. and R.L. co-supervised the project. F.L., R.L., and T.L. designed experiments. B.J.P. performed Pol II CTD mutational analyses and contributed to crucial preliminary data. R.F. and J.E.W. designed and generated the *Drosophila* IntS8 antibody. D.S.G. generated the Halo-Rpb1 fly line and supported the initial phase of the project. F.L. performed all other experiments. F.L. and T.L. analyzed the data. F.L., R.L., and T.L. interpreted the data. F.L. drafted the manuscript. F.L. T.L., R.L., D.S.G., J.E.W., and B.J.P. revised the manuscript.

## Data, code, and materials availability

Sequencing data have been deposited to the Gene Expression Omnibus (GEO) under the accession no. GSE33499. Table S1 and S2 lists all the HCR FISH probes used in this study. All fly lines used in this study are available upon request. Data analysis scripts are available through Github https://github.com/timotheelionnet/Pol2ClustersEggChamber.

## Supplementary Materials

Materials and Methods

Figs. S1 to S12

Tables S1 to S2

References (*1–98*)

Movies S1 to S2

