## Supplement for "Transcription machinery clustering by Integrator synchronizes histone gene expression"

### Materials and Methods

#### *Drosophila* strains

CRISPRed Halo-Rpb1 fly line was generated through the injection of DNA mixtures into a Cas9-expressing fly line (Bloomington 51324). As previously described(*57*), each DNA mixture contains two CRISPR targeting vectors that express sgRNAs and a homology donor vector. The two CRISPR targets were selected from a list of targets with zero predicted potential off-target sites using flyCRISPR Optimal Target Finder (http://tools.flycrispr.molbio.wisc.edu/targetFinder/, Maximum Stringency, NGG PAM sequence only). Complementary strands encoding the sgRNAs were annealed, phosphorylated, and cloned into the BbsI site of pU6-BbsI-chiRNA (Addgene plasmid 45946). The combination of Rpb1_fw1: 5'-CTTCGTAGGGATTTGAGAGCCAGTG-3' and Rpb1_rev1: 5'-AAACCACTGGCTCTCAAATCCCTAC-3' directed cleavage downstream of the 3'UTR of Rpb1.The combination of Rpb1_fw2: 5'-CTTCGAGAAACACTCGGCGAGGCT-3' and Rpb1_rev2: 5'-AAACAGCCTCGCCGAGTGTTTCTC-3' directed cleavage adjacent to the start of the *Drosophila* CTD. Two homology arms, each about 1 kb in length, were PCR amplified from strain $w^{1118}$. The homology arm adjacent to the 3'UTR of Rpb1 was cloned into the AarI site of pHD-ScarlessDsRed (DGRC plasmid 1364). The other homology arm was cloned into the SapI site alongside desired mutations in the CTD and the CRISPR targeting site. Candidate recombinant flies were identified by expression of the DsRed marker in eyes. Individual DsRed candidates were out-crossed to strain $w^{1118}$. Desired candidates were verified by sequencing and western blot. The *Halo-Rpb1* transgene was combined with *GFP-Mxc*(*42*) (Bloomington 92330) or *GFP-CycE*(*37*) through genetic matings.

All RNAi experiments involving NCs were driven by a *Mat-GAL4* fly line that carries two genomic insertions of the *Mat-GAL4* transgene, one on the second chromosome (Bloomington 7062) and the other on the third chromosome (Bloomington 7063). *UAS-mCherry-RNAi* (Bloomington 35785) was used as a control. Other RNAi fly lines used in the study: *UAS-IntS8-RNAi* (Bloomington 33048). *UAS-IntS11-RNAi* (Bloomington 65093). *UAS-IntS9-RNAi* (Bloomington 65892). The *UAS-IntS9-RNAi* fly line from the Bloomington stock center is heterozygous and was outcrossed to a *CyO* balancer strain to generate a homozygous line.

All stocks and crosses were maintained at 22°C on Molasses medium.

#### *Ex-vivo* ovary culture for live imaging and chemical treatments

Ovaries from well-fed females were dissected in growth media consisting of Schneider's medium (Fisher 21720-001) supplemented with 10% fetal bovine serum (Invitrogen 10082-147), 0.2mg/ml bovine insulin (Sigma-Aldrich I0516), and 0.6x streptomycin/penicillin (Thermo Scientific 15140122)(*82*). To visualize Halo-tagged factors, the ovaries were incubated in growth media containing 100 nM of either Janelia Fluor 549 (JF549) or Janelia Fluor 646 (JF646)(*80*) for 1 hour on a horizontal shaker. Ovaries were then transferred to a MatTek dish (MatTek Corporation P35G-1.5-20-C) and embedded in 0.8% low-melting temperature agarose (NuSieveGTG from Lonza)(*83*) in the presence of ~3 ml of surrounding growth media. For chemical treatments, agarose-embedded ovaries were incubated in growth media containing 500 nM FP (Santa Cruz Biotechnology, sc-202157; in DMSO) and 500 μM TRI (TOCRIS Bioscience, 3253; in DMSO)(*52*), respectively. Ovaries incubated in growth media containing the same concentrations of DMSO for the same incubation times were used as controls.

Immunofluorescence

For polytene chromosome staining, salivary glands from wandering third-instar larvae were dissected and incubated in Schneider's medium containing 100 nM JF646 for 50 minutes. For heat shock, the larvae were placed in a 37°C water bath for 20 minutes before dissection. The salivary glands were subsequently squashed as previously described (Schwartz et al., 2004). Slides were incubated with mouse ARNA3 antibody (EMD Millipore CBL221, 1:200) overnight at 4°C, then probed with goat anti-mouse IgG (1:500; Alexa Fluor 488) for 4 hours at room temperature. Samples were imaged on a Zeiss LSM 800 laser scanning confocal microscope and adjusted for brightness and contrast using ImageJ.

Ovaries were dissected in Schneider's medium, fixed in 4% PFA (EMS 15713) with PBS+0.5% Triton (PBST) for 20 minutes at room temperature, and washed three times for 10 min with PBST. Ovaries were blocked for 1 hour in 0.2% BSA in PBST (Thermo Scientific AM2618) at room temperature, then incubated with primary antibodies overnight at 4°C. Ovaries were washed three times for 1 hour with PBST before incubation with secondary antibodies for three days at 4°C. Ovaries were washed three times for 1 hour with PBST, and once for 1 hour with PBS before incubation with VECTASHIELD (Vector Laboratories H-1000-10) overnight at 4°C. Hoechst was added to the second of the final three PBST washes to visualize DNA. Primary antibodies: mouse ARNA3 (EMD Millipore CBL221, 1:200), rat anti-Ser2ph (Sigma-Aldrich 04-1571, 1:200), rat anti-Ser5ph (EMD Millipore 04-1572, 1:200), rabbit anti-Spt5 (1:500), rabbit anti-CycT (gift from Akira Nakamura(*84*), 1:500), mouse MPM-2 (Sigma-Aldrich 16-220, 1:2000), guinea pig anti-Mxc (gift from Robert Duronio, 1:2000), rabbit anti-HA (Thermo Fisher Scientific 715500, 1:200), mouse anti-FLAG (Sigma-Aldrich F1804, 1:200), rabbit anti-IntS8 (*85*) (1:500). For experiments involving JF dyes, ovaries were first incubated in 100 nM JF dye containing growth media consisting of Schneider's medium supplemented with 10% fetal bovine serum, 0.2mg/ml bovine insulin, and 0.6x streptomycin/penicillin for 1 hour prior to fixation. For combined detection of MPM-2 and RNA, a Cy5 conjugated-MPM-2 antibody (Sigma-Aldrich 16-220) was used to shorten incubation times. Secondary antibodies were Alexa Fluor 488, Cy3, and 647 (Jackson ImmunoResearch) at 1:500.

Unless otherwise noted, all live and fixed imaging experiments were performed on a Nikon W1 spinning disk confocal microscope with an SR HP Plan Apo 100× 1.35 NA objective and Andor 888 electron-multiplying charge-coupled device (EMCCD) cameras driven by Nikon Elements software.

Western blotting

5 pairs of ovaries were dissected in Schneider's medium, homogenized, and boiled in LDS sample buffer (Invitrogen). Tissue lysates equivalent to 0.25 ovary were loaded into each lane on a 3–8% Tris-acetate SDS–polyacrylamide gel electrophoresis (PAGE) gel (Life Technologies). Halo-tagged fusion proteins were detected using rabbit anti-Halo antibody (Promega G9281, 1:2000), Rpb1 was detected using mouse ARNA3 antibody (EMD Millipore CBL221, 1:3000).

Spt5 was detected with rabbit anti-Spt5 antibody (1:3000)(*86*). Blots were subsequently probed with fluorescently labeled goat anti-rabbit IgG (1:3000; Alexa Fluor 488) and goat anti-mouse IgG (1:3000; Alexa Fluor 647) and visualized with a BioRad scanner.

CUT&RUN library preparation

CUT&RUN libraries were prepared using the CUTANA™ CUT&RUN Kit (EpiCypher, Cat. No. 14-1048) following the manufacturer's instructions with minor modifications. In brief, ovaries were harvested from 0-6 hr old adult females fattened with yeast paste for 8-10 hrs to reach development Stage 10. Four pairs of ovaries were collected per replicate, with two to four replicates prepared per genotype per condition. For drug-treated conditions, ovaries were incubated in growth media containing either 500 µM TRI or 500 nM FP for 35 min. To allow for antibody binding in situ, cells were permeabilized in buffer containing 0.05% digitonin. The ovaries were incubated with rabbit polyclonal Halotag antibody (Promega G9281; 1:100) at 4°C overnight. The $w^{1118}$ line, lacking HaloTag expression, served as a negative control. Following antibody incubation, pAG-MNase (EpiCypher, Cat. No. 15-1016) was added to bind the antibody-labeled chromatin. Targeted digestion was activated by the addition of 100 mM $CaCl_2$. To normalize read counts across conditions, 0.1 ng of E. coli Spike-in DNA (EpiCypher, Cat. No. 18-1401) was added to each reaction within the Stop Buffer master mix used to halt MNase activity.

CUT&RUN-enriched DNA was isolated from the supernatant and purified using 1.66x SPRIselect beads (Beckman Coulter, Cat. No. 21-1405). Purified DNA was quantified via Qubit fluorometer. Sequencing libraries were generated using universal i5 and uniquely barcoded i7 primers with 14 PCR cycles. All subsequent library purifications were performed using 1.66x SPRIselect beads to eliminate residual primers and concentrate the samples. Final library quality and fragment distribution—targeting a peak at ~300 bp representing mononucleosomes plus sequencing adapters—were confirmed via Bioanalyzer prior to paired-end sequencing on an Illumina NextSeq 550.

CUT&RUN data processing

Paired-end CUT&RUN sequencing reads were assessed with FastQC (*87*) using default criteria, then adapter and quality trimming were performed using Trim Galore (*88*) in paired-end mode. Trimmed reads were aligned with Bowtie2 (*89*)(local, very-sensitive-local) to an *E. coli* reference genome to quantify spike-in reads and to a custom *Drosophila melanogaster* genome: starting from the dm6 assembly, we swapped the histone locus repeats for a single instance of the consensus sequence of the ~ 5kb repeated unit, followed by a stretch of unspecified bases to retain the register of Chr. 2L regions downstream of the histone locus with the dm6 assembly. This procedure prevents the undesirable exclusion of histone locus reads that do not map unambiguously to a unique repeat. As a result, reads from the ~110 histone repeats pile up on a single unit, ensuring ample read depth and providing an average profile of the repeat unit. Alignments were processed with SAMtools (*90*), including sorting/indexing and filtering by a minimum mapping quality, and reads mapping to the mitochondrial genome were removed. Spike-in normalization was applied by computing a per-sample scaling factor from the number of mapped *E. coli* reads, and scaled genome-wide coverage tracks were generated using deepTools (*91*) bamCoverage. Replicates were then averaged to generate a mean bigWig track, and background-corrected bigWigs were produced by subtracting from each coverage track the coverage track obtained from non-Halo-expressing negative-control ovaries submitted to the same biological treatment and processed identically. Signal around transcription start sites was summarized with deepTools computeMatrix and visualized using plotHeatmap and plotProfile.

For gene-level quantification, CUT&RUN signal was first summarized per gene and per sample as a single numeric value (summed signal across the gene), and these values were then aggregated across all samples for the genes of interest. For each gene, background signal was estimated separately for each treatment using the matched negative-control samples and averaged across replicates, then subtracted from the corresponding experimental samples to obtain background-corrected gene signals. To calculate the summary matrix used for plotting, the background-corrected control condition for each comparison was scaled to 1 by dividing both the control and treatment background-corrected signals by the mean background-corrected control signal computed across replicates for that gene. Results were then visualized as replicate-level points with the mean indicated for each condition.

Quantification of HLB enrichment from live imaging movies

To quantify the recruitment of transcription regulators to HLBs in live movies, sum-intensity projections were generated using five consecutive z-slices centered on the focal plane of the clusters. ROIs were defined by segmenting the HLB using the channel with the higher signal-to-noise ratio. We computed the average intensity in each ROI, and background-corrected the result by subtracting the mean nucleolar intensities. To estimate enrichment values before the first appearance of each cluster, we measured the average intensity at the ROI where we first observed a detectable signal accumulation at the two preceding time points. To ensure tracking fidelity, any nurse cells (NCs) characterized by significant xyz translational movement were excluded from the quantification.

FRAP experiments

Ovaries from females expressing both CycE-GFP and Halo-tagged factors were dissected in growth media and subsequently incubated for 1 hour in growth media containing 100 nM of Janelia Fluor 552 (JF552)(*81*). For drug-treated samples, ovaries were preincubated in growth media containing the indicated drug for 35 min before FRAP imaging. To minimize egg chamber movement, the stained ovaries were further dissected into single ovarioles, and egg chambers beyond Stage 10 were removed before being transferred onto a concanavalin A (50 µg/ml, Cayman Chemical 14951)–coated MatTek dish. Ovaries were then embedded in 0.8% low-melting temperature agarose in the presence of ~1 ml of surrounding growth media.

FRAP experiments were performed on a Zeiss LSM 880 laser scanning confocal microscope with an LD C-Apochromat 40x/1.1 W Korr M27 objective. Four images were taken before bleaching. In the fifth frame, a circular region surrounding the cluster of interest was bleached with full laser power, and recovery was recorded in a single z-plane at 1% power at 1s intervals for at least 300s. FRAP movies with noticeable z-drift are excluded from the analysis; xy-drift correction was performed using the FIJI Template Matching plugin (https://sites.google.com/site/qingzongtseng/template-matching-ij-plugin?authuser=0). Normalized fluorescence intensity was calculated with a three-step normalization: first by subtracting from all intensity values the average signal of a region outside of nuclei to obtain background-subtracted values, second by normalizing the background-subtracted fluorescence intensity of Halo fusion to that of an unbleached region to correct for the subtle loss in fluorescence intensity over imaging times, and third by normalizing this ratio to that of the pre-bleach spot. The recovered fractions were calculated by averaging the last 11 points on the recovery curves. Half-recovery times were estimated by interpolating the FRAP curves at the half-recovered fraction.

Fitting of FRAP recovery curves

For fig. S6A, FRAP recovery curves were analyzed in Python by fitting the normalized intensity time courses to multi-exponential recovery models using nonlinear least-squares regression. As a preliminary step, the rate constant for the fast-recovery component was estimated by fitting nucleoplasmic recovery traces (recorded outside the HLB) with a multicomponent exponential model; this value subsequently constrained the minimum rate allowed for the fastest component in subsequent FRAP fits. Each dataset was fitted to both a double-exponential model, $f(t) = 1 + A1\ e\text{-}a1t + A2\ e\text{-}a2t$ , and a triple-exponential model, $f(t) = 1 + A1\ e\text{-}a1t + A2\ e\text{-}a2t + A3\ e\text{-}a3t$ , using bounded parameters and requiring that at least one component represent a fast recovery phase. To mitigate overfitting in the triple-exponential model, additional rejection criteria were applied to exclude degenerate solutions characterized by overlapping rate constants, excessively slow components, or insufficient separation between rates. Fit quality was evaluated using $R^2$ and residual sums of squares, and model selection was performed using Akaike and Bayesian information criteria (AIC and BIC) (*92*, *93*) computed under a Gaussian error assumption; the preferred model was identified by lower information criteria values. Best-fit parameters were ranked by rate to classify components by recovery speed, and visual inspection of residuals was used to validate model alignment.

The analysis in fig. S6B was performed by jointly fitting averaged recovery curves from the Pol II and Spt5 datasets of matching conditions to a shared three-component exponential model. The rationale for this procedure is that Pol II and Spt5 form a stable complex during pausing and elongation (*94*), and thus the kinetic rate corresponding to this state is expected to be identical for both factors. Replicate recovery traces were first interpolated onto a common time grid and averaged; these resulting mean curves were then simultaneously fitted by nonlinear least-squares regression. The model allowed factor-specific values for all kinetic component amplitudes and two of three rate constants, while enforcing a single shared rate constant between the two datasets. Fast-component rates were fixed to pre-determined values derived from the nucleoplasmic fits, while the remaining parameters were estimated within bounded constraints. Fit quality was assessed independently for each condition, and kinetic parameters were reported as rate constants and corresponding half-times for the fitted components.

Antibody production

The N-terminal region of *Drosophila* IntS8 (amino acids 1-308) was cloned into pRSFDuet-1 using BamHI and HindIII restriction sites to generate pRSFDuet-1 IntS8 AA 1-308 (Addgene #196905). The plasmid was then transformed into BL21 Star (DE3) *E. coli* and grown in terrific broth media supplemented with 50 µg/mL kanamycin. Expression of His-tagged IntS8 was induced at OD600 ~0.8 by addition of 0.3 mM IPTG and cells were incubated at 25°C for 7 h before harvesting. His-tagged IntS8 was purified under denaturing conditions and all purification steps were carried out at 4°C unless otherwise noted. Cells were resuspended and incubated in lysis buffer (30 mM Tris pH 7.5, 300 mM NaCl, 1 mM DTT, 1 mM EDTA, 100 µg/mL lysozyme, 1 mM PMSF, 100 µM leupeptin, 10 µM pepstatin A, and 1 mM benzamidine) on ice for 30 min, and then sonicated for a total of 5 min. Sample was pelleted by centrifugation at 20,000 x g for 15 min and then resuspended in denaturing buffer (30 mM Tris pH 7.5, 300 mM NaCl, and 8 M urea) at 4°C for 30 min while stirring. Cell debris was removed by centrifugation at 20,000 x g for 15 min. The solubilized IntS8 protein was diluted with buffer to lower the urea concentration to 1 M, loaded onto a Ni-column (Cytiva 29051021), and eluted by adding elution buffer (30 mM Tris pH 7.5, 300 mM NaCl, and 500 mM imidazole). Fractions containing His-tagged IntS8 were pooled, dialyzed against PBS, concentrated, flash frozen, and stored at -80°C. Purified protein in PBS was shipped to Cocalico Biologicals and used to inoculate rabbits. The reactivity and specificity of antisera was confirmed with Western blots using whole cell extracts from Drosophila DL1 cells treated for 3 d with dsRNAs targeting IntS8.

EdU labeling

Ovaries were dissected in growth medium and incubated with 10 µM EdU for 15 min. EdU-treated ovaries were then rapidly rinsed three times and cultured in EdU-free growth medium for 75 min before fixation in 4% PFA in PBST for 20 min, followed by two 5-min washes in PBST. Incorporated EdU was detected by click chemistry using Alexa Fluor 488 azide according to the manufacturer's instructions (Thermo Fisher Scientific C10337).

RNA in situ hybridization

All steps were performed using RNAse-free reagents following a procedure adapted from previous protocols(*95, 96*). Ovaries were dissected in growth medium, fixed in 2% PFA in PBST at room temperature for 55 min, and washed three times with PBST for 10 min. The fixed ovaries were dehydrated with graded EtOH series: 30%, 50%, 70%, 100%, 100%, 100%, 10 min each and incubated at 4°C overnight on a shaker. The dehydrated ovaries were rehydrated with graded EtOH series: 70%, 50%, 30%, 10 min each, then incubated at 4°C 5% acetic acid for 5 min. The ovaries were washed three times with ice-cold PBS for 5 min and post-fixed in 2% PFA at room temperature for 55 min. Ovaries were pre-hybridized in probe hybridization buffer (Molecular Instruments) for 30 min at 37°C, then incubated with the desired probe sets (see Tables. S1 and S2) at 37°C for 20 hours. Samples were washed four times with probe wash buffer (Molecular Instruments) for 15 min at 37°C and twice with 5 X SSCT (5 X sodium chloride sodium citrate, 0.1% Tween 20) at room temperature. Ovaries were incubated in the amplification buffer (Molecular Instruments) for 15 min at room temperature. The amplification buffer was subsequently removed and the hairpin solution (prepared by heating 6 pmol of each hairpin (Molecular Instruments) for 90 sec at 95°C, cooling at room temperature in the dark for 30 min, and subsequently adding the snap-cooled hairpins to the amplification buffer at room temperature) was added to each sample. Samples were incubated for 20 hours in the dark at room temperature, and washed five times with 5 X SSCT, twice for 5 min, twice for 30 min, and once for 5 min. Hoechst was added to the first 30 min wash for DNA detection. For combined MPM-2 antibody detection, ovaries were first fixed in 2% PFA in PBST at room temperature for 55 min, and washed three times with PBST for 10 min, then proceeded with the immunofluorescence procedure from the blocking step. After the final antibody wash, antibody-stained ovaries were subsequently dehydrated with the graded EtOH series and processed through the HCR FISH procedure.

The DS-H3 probe set was synthesized as an Oligo Pools (Table S2, Integrated DNA Technologies). The rest of the probes were ordered as individual oligonucleotides (Table S1, Integrated DNA Technologies).

Semi-automated NC nuclei segmentation and cluster detection

A Fiji-based(*97*) semi-automated analysis pipeline was developed to detect nuclei, clusters and quantify their intensity based on immunofluorescence confocal z-stacks of NCs (all Fiji macros and Matlab scripts available at https://github.com/timotheelionnet/Pol2ClustersEggChamber). In brief, images containing individual egg chambers of desired stages were first detrended along the Z axis to correct for intensity losses with increasing tissue depth. Nuclei within the detrended images were initially segmented based on the Hoechst channel using the segmentNuclei_v2 macro with the following parameters: AverageNucleiDiameter = 20 µm, MinVolume = 400 $\mu m^3$, MaxVolume = 40,000 $\mu m^3$, MaxSurfaceToVolumeRatio=1, MinSphericity=0.4. Briefly, image stacks were subjected to a bandpass filter followed by intensity thresholding; small objects were eliminated using a morphological opening, followed by a watershed step in order to split conjoined nuclei. Following nuclei segmentation, clusters were detected using the clusterAnalysisFromSegmentedNuclei_v2 macro with the following parameters: Radius of Median Filter to smooth cluster signal = 2, Radius of Median Filter to average out background = 50, Threshold Factor to call clusters (multiples of nucleoplasm StdDev) = 3. First, a mask marking the nucleoplasm (i.e., excluding nucleoli) was generated within each nucleus by thresholding the nuclear factor intensity using the Otsu algorithm. Transcription clusters were then detected using either the Ser5ph (if available) or the factor of interest (when the Ser5ph channel is unavailable) channels. At each Z-slice, the nucleoplasm background intensity (average signal across the nucleoplasm mask) was subtracted; following a median filtering step, clusters were detected using a threshold equal to three times the standard deviation of the background-corrected nucleoplasm intensity, and their intensity metrics were measured. Following cluster detection, subsequent analysis steps were performed in Matlab using custom scripts. Clusters were assigned as HLBs if their volume was 1 $\mu m^3$ or more - this minimal value was based on the observation that all HLBs in this size range were Mxc positive. In each nucleus, the fluorescence background signal was subtracted from all cluster intensity values. The background was computed as the average signal of the corresponding channel across the nucleoli masks of the same nucleus. When normalizing for nuclear expression levels, we estimated the nuclear intensity as the average intensity across the nucleoplasm mask, subtracted by the background intensity (similarly computed as the average signal of the corresponding channel across the nucleoli masks). NCs with fewer than three clusters above 1 $\mu m^3$ are typically false positive HLB detections based on their morphology distinct from that of *bona fide* HLBs and were discarded. For analyses involving the cell cycle stages, HLBs whose average MPM-2 or P-TEFb intensities were more than (X+0.5) times the average nucleoplasmic intensities were classified as S-phase NCs, whereas HLBs whose average MPM-2 or P-TEFb intensities were less than (X-0.5) times were classified as G-phase NCs. X corresponds to the minimally detectable enrichment level and was determined empirically for each dataset by sampling NCs whose MPM-2 or P-TEFb intensities at HLBs were barely above nucleoplasmic intensities by eye. NCs whose averaged HLBs were X±0.5 were manually examined for their cell cycle stage classification.

RNA FISH quantification

For quantifications in Fig. 6B, fig. S10B, and fig. S12B, H3 RNA FISH was quantified using FIJI by measuring the average signal intensity in a manually outlined cytoplasmic region of each NC. For quantifications in Fig. 6E and fig. S10D, RpL30 RNA FISH was quantified using FIJI by measuring the average signal intensity in a manually outlined cytoplasmic region of each individual egg chamber. For Fig. 6C and fig. S10C, nuclei containing MPM-2 enriched foci consistent in size and morphology with HLBs were scored as S-phase; other nuclei were scored as G-phase. NCs with H3 RNA FISH intensities measured in the manually outlined cytoplasmic region above 2.1 SD from the mean intensities in the mCherry-RNAi control condition were scored as H3-positive. For fig. S11C, NCs whose 3'-H3 intensities at HLBs and in the cytoplasmic particles were both above 1,500 AU were chosen for the analyses to improve measurement reliability by minimizing the impact of background signal. A Maximum Intensity Projection of 25 slices centered around the NCs of interest (z-stack step size = 0.3 µm) was generated, and the background intensities were quantified by measuring the average signal intensity in manually outlined regions in the neighboring G-phase NCs. Each particle intensity was quantified by measuring the maximum intensity in a manually outlined region marking each individual focus. The final DS-H3 to 3'-H3 ratios were calculated by the division of their background-corrected maximum intensity values.

Statistical Significance tests

Unless otherwise noted, all statistical significance tests were performed using two-tailed, unpaired t-tests with Welch's correction. P-values greater than 0.05 were considered to indicate that the conditions compared were not statistically significantly different.

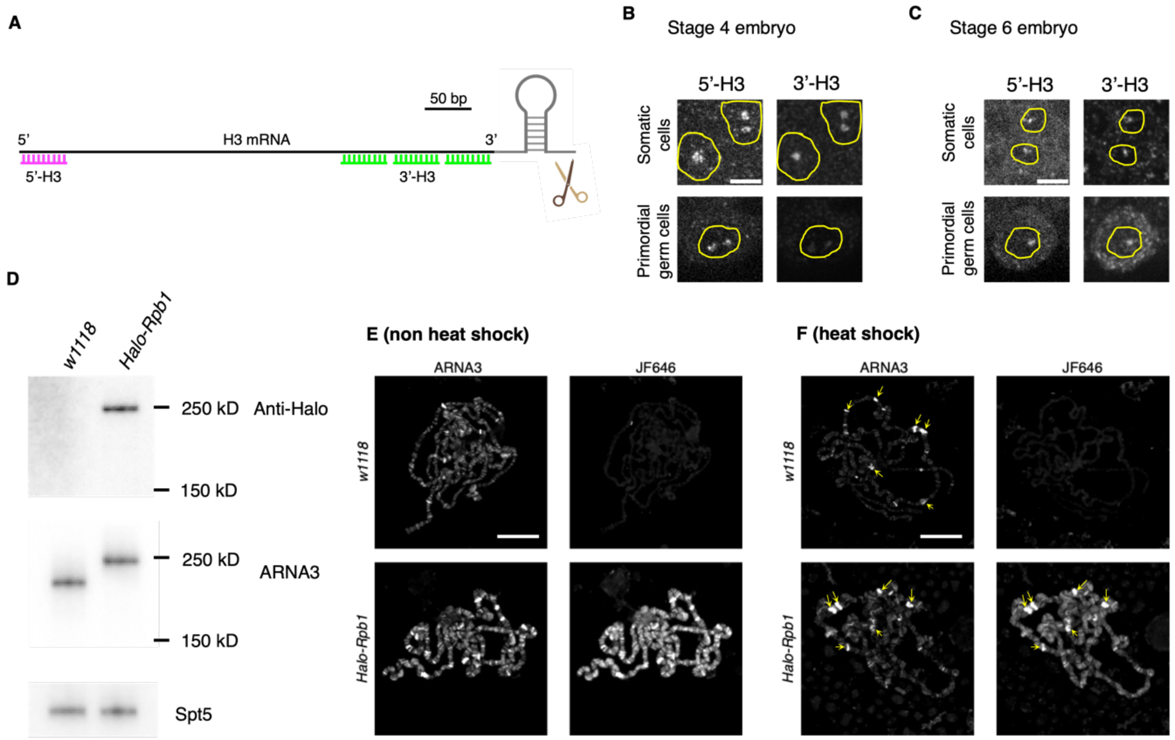

**Fig. S1. Validation of the H3 FISH probes and CRISPR-ed Halo-Rpb1 fly line.** Related to **Fig. 1**. (**A**) Schematic of the 5′ (magenta) and 3′ (green) H3 probe design (drawn to scale). The 5' probe targets the first ~50 bp and thus is expected to hybridize to the nascent transcript produced by Pol II before the promoter-proximal pause. Fully transcribed pre-mRNAs and mature mRNAs should hybridize to both 5' and 3' probes equally. Scissors indicate the location of the 3' cleavage site. (**B** and **C**) Validation of histone FISH probes in somatic cells and primordial germ cells (PGCs) of embryos. The 5′ H3 probe produces distinct nuclear FISH foci in both somatic cells and PGCs across embryonic stages. The 3′ H3 probe produces distinct nuclear foci in somatic cells, but in PGCs, it yields clear nuclear foci only at later stages (C), coincident with the release of Pol II from the promoter proximal pause (*51*, *84*) in later, but not earlier stages (A). Nuclei are outlined by the yellow circles. Scale bars, 5 µm. (**D**) Western blot detection of the Halo-Rpb1 using anti-Halo and ARNA3 (against Rpb1) antibodies, respectively, in adult ovary lysates. $w^{1118}$ was used as a negative control. Spt5 serves as a loading control. (**E** and **F**) Detection of Halo-Rpb1 on fixed and spread polytene chromosomes in the larval salivary glands before (E) and after (F) heat shock. Pol II is redistributed to the major heat shock puffs (marked by yellow arrows) upon heat shock. ARNA3 detects both Halo-tagged and unmodified versions of Rpb1, whereas JF646 specifically detects the Halo-tagged Rpb1. Scale bars, 50 µm.

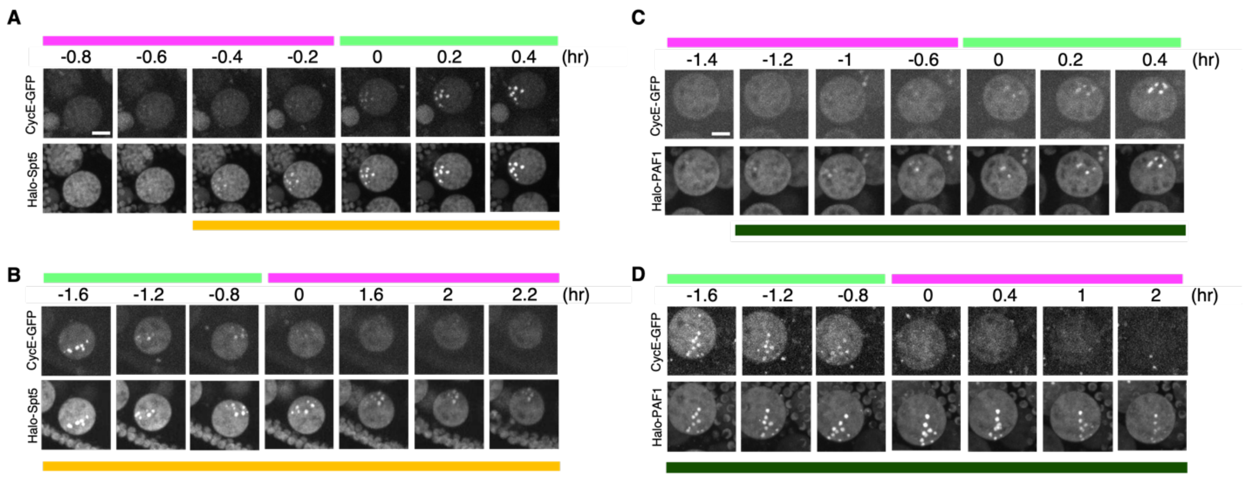

**Fig. S2. Representative frames from time-lapse movies of nuclei entering or exiting S phase.** Related to **Fig. 1**. Representative time-lapse snapshots of Halo-Spt5 (A,B), and Halo-PAF1 (C,D) in nuclei entering (A,C) or exiting (B,D) S phase. S-phase nuclei are identified by enrichment of CycE–GFP at the HLBs. Time 0 denotes the onset of S phase (entering panels) or the onset of G phase (exiting panels). Halo-tagged proteins were labeled with Janelia Fluor 549. Top magenta bar marks G phase frames, top light green bar marks S phase frames; bottom color bar marks frames where the factor is detected at the HLB. Scale bar, 10 µm.

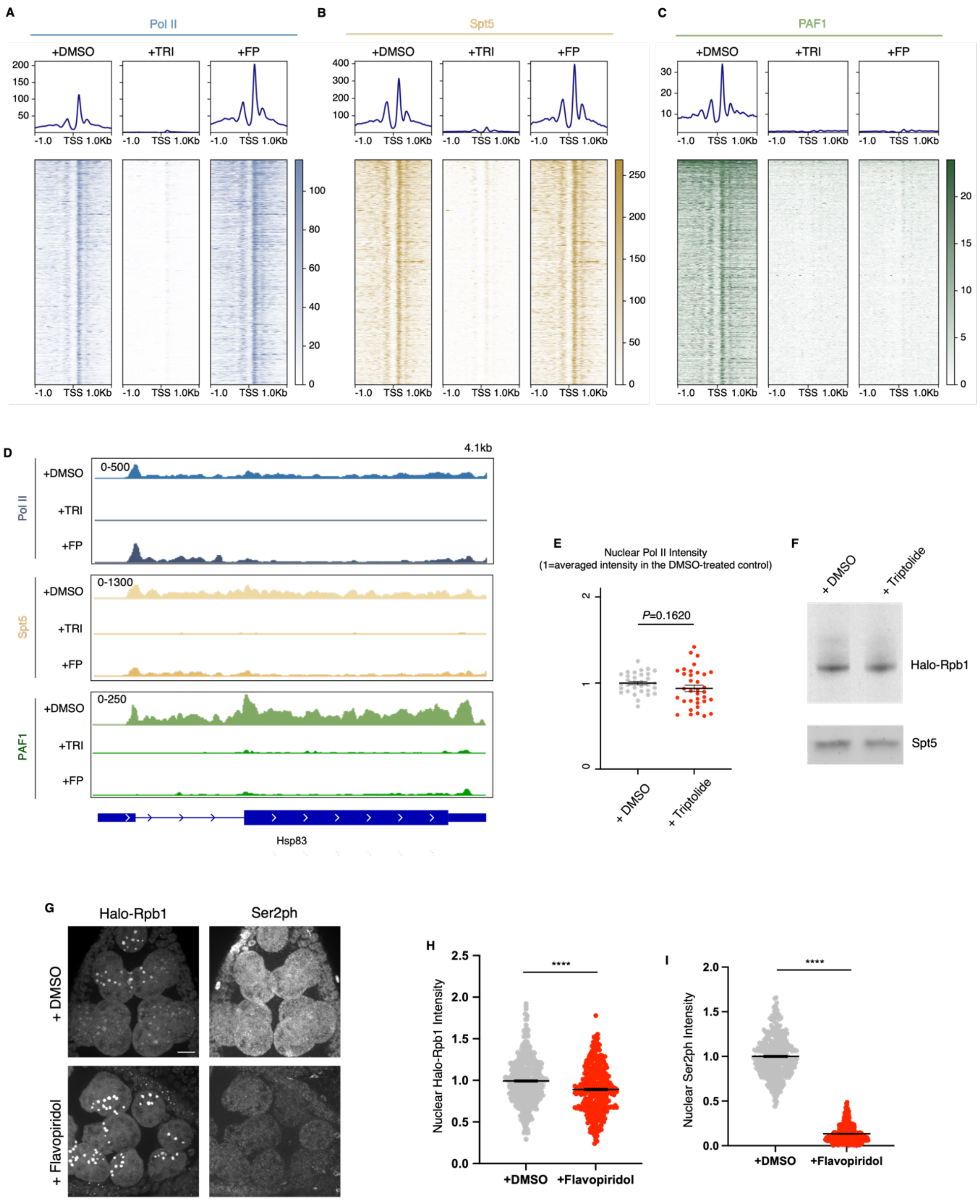

**Fig. S3. Validation of the effects of triptolide (TRI) and flavopiridol (FP) on transcription.** Related to **Fig. 2**. (**A** to **C**) Promoter enrichment profiles of Pol II (A), Spt5 (B), and PAF1 (C). Heatmaps (bottom) and metaplots (top) for Pol II (blue), Spt5 (orange), and PAF1 (teal) measured by Halo CUT&RUN at the 5,000 promoters with highest occupancy for the respective factor. Signals are centered on the transcription start site (TSS) ± 1 kb; promoters are ordered by decreasing occupancy in the DMSO control. Read counts were spike-in calibrated, subtracted by similarly calibrated read counts from the non-Halo expressing control samples (submitted to matching perturbations and antibody incubations), and represent the mean of at least two biological replicates. (**D**) Pol II, Spt5, and PAF1 CUT&RUN occupancy at *hsp83* in DMSO control, TRI- and FP-treated conditions. TRI treatment leads to a near-complete loss of Pol II, Spt5, and PAF1 signal across the entire locus, consistent with inhibition of transcription initiation (*52, 53*). In contrast, FP treatment depletes Pol II, Spt5, and PAF1 signals from the gene body, while Pol II and Spt5 remain enriched near the TSS, consistent with inhibition of pause release (*52, 53*). (**E** and **F**) Halo-Rpb1 is not degraded upon 35 min of 500 μM Triptolide treatment. (E) Quantifications of the nuclear Pol II intensity in DMSO-treated versus TRI-treated Halo-Rpb1 expressing egg chambers. Each dot represents the averaged nuclear Pol II intensity in one egg chamber. Error bars represent mean ± SEM. Data were acquired from 33 DMSO-treated and 36 TRI-treated stage-matched egg chambers derived from 10 females per condition. (F) Western Blot detection of Halo-Rpb1 in DMSO-treated versus TRI-treated egg chambers using anti-Halo antibody. Spt5 serves as a loading control. (**G** to **I**) FP reduces Pol II Ser2 phosphorylation. Representative confocal images of *Drosophila* egg chambers treated with DMSO or 500 nM FP for 35 min (G). Scale bar, 5 μm. Quantification of nuclear intensity for Halo-Rpb1 (H) and Ser2ph (I). Total Pol II (Halo-Rpb1) is largely unchanged, whereas Ser2ph is strongly reduced upon FP treatment. Each dot represents one nucleus; error bars indicate mean ± SEM from 6 stage-matched egg chambers per condition. ****$P < 0.0001$ (two-tailed Student's t-test).

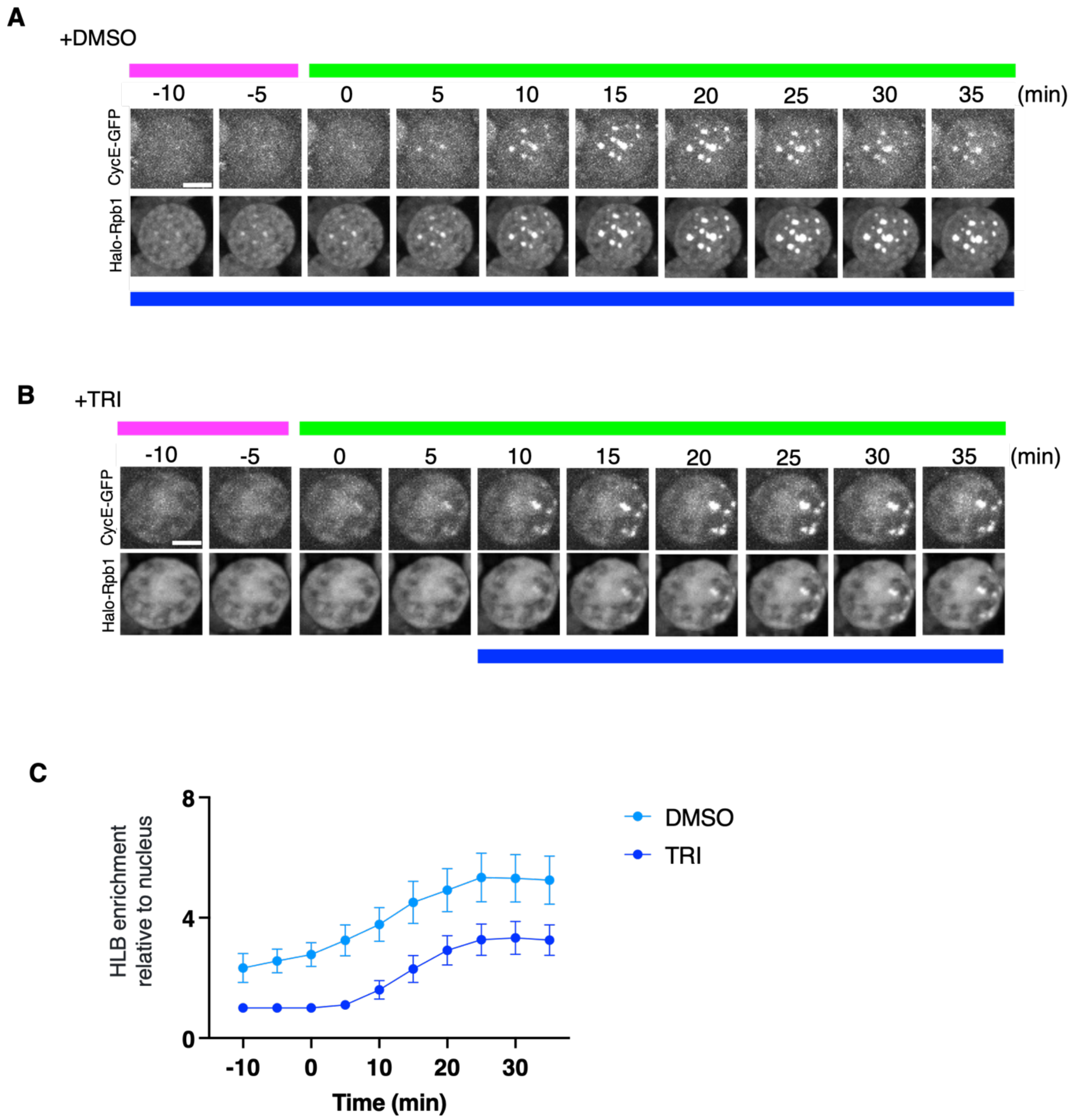

**Fig. S4. Pol II clusters form at the HLBs after S-phase entry even when transcription initiation is inhibited.** Related to **Fig. 2**. (**A** and **B**) Representative frames from time-lapse movies of nurse cell nuclei entering S phase in control egg chambers treated with DMSO (A) or in egg chambers treated with TRI (B). Halo–Rpb1 was labeled with Janelia Fluor 549. Time 0 denotes S-phase entry. Top magenta bar marks G phase frames, top light green bar marks S phase frames; bottom color bar marks frames where Pol II is detected at the HLB. Scale bar, 10 µm. (**C**) Fluorescence enrichment at HLB relative to the nucleoplasm average for Halo-Rpb1 at HLBs as NCs enter S-phase. *n* = 3 egg chambers per condition. Error bars, mean ± SEM.

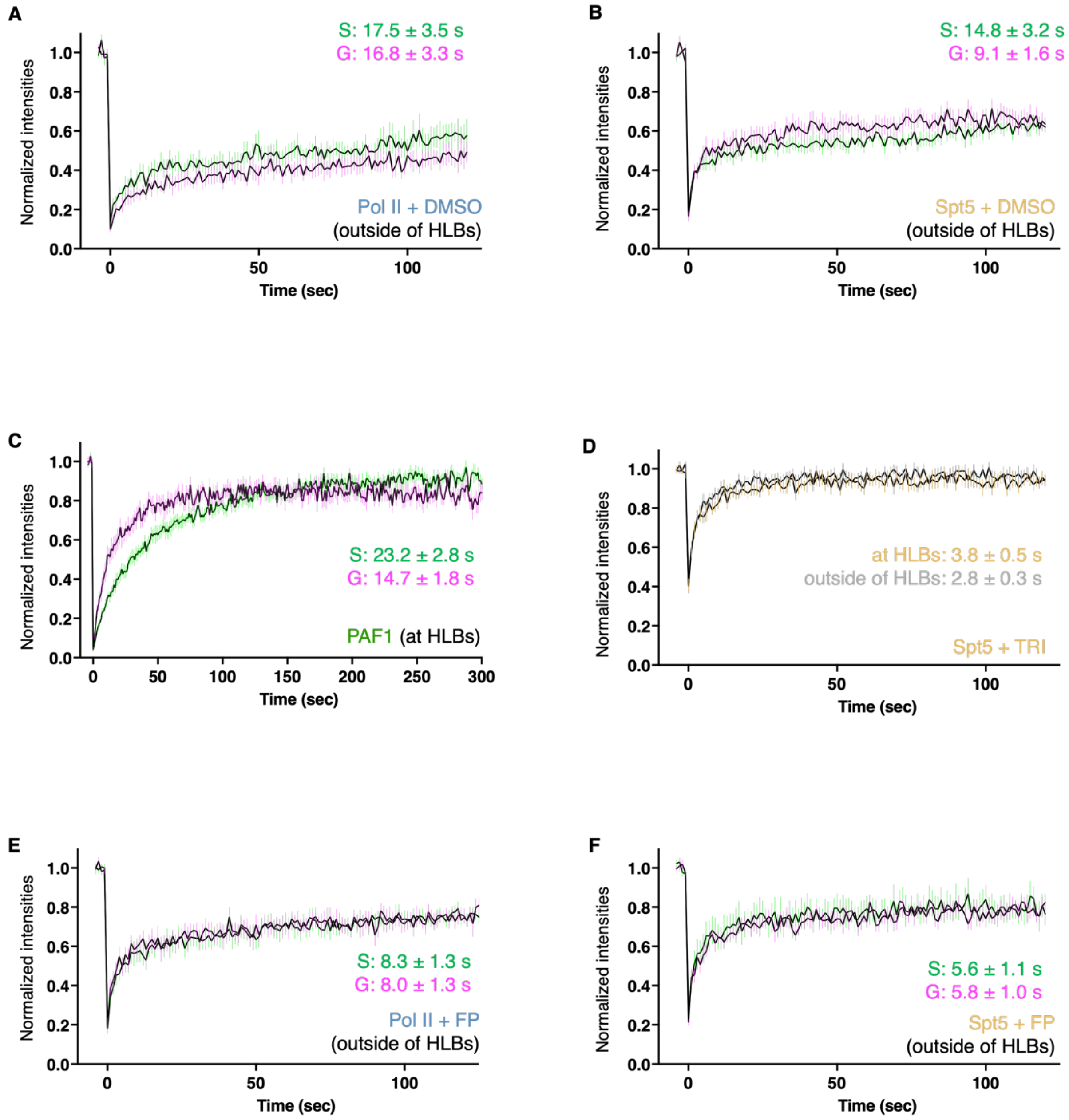

**Fig. S5. Additional FRAP analysis of Pol II, Spt5, and PAF1.** Related to **Fig. 3**. (**A** and **B**) FRAP curves of G-phase (magenta) and S-phase (green) HLB-bound Pol II (A) and Spt5 (B) clusters outside HLBs in DMSO control treatment. (**C**) FRAP curves of G-phase (magenta) and S-phase (green) HLB-bound PAF1 clusters at HLBs. (**D**) FRAP curves of Spt5 at (orange) versus outside of (gray) HLBs in TRI treatment. (**E** and **F**) FRAP curves of G-phase (magenta) and S-phase (green) HLB-bound Pol II (E) and Spt5 (F) clusters outside HLBs in FP treatment. Black lines: average curves from 7-10 nuclei across at least 5 ovaries. Vertical colored lines, SEM. A single exponential recovery curve was fitted to each factor to estimate half-lives.

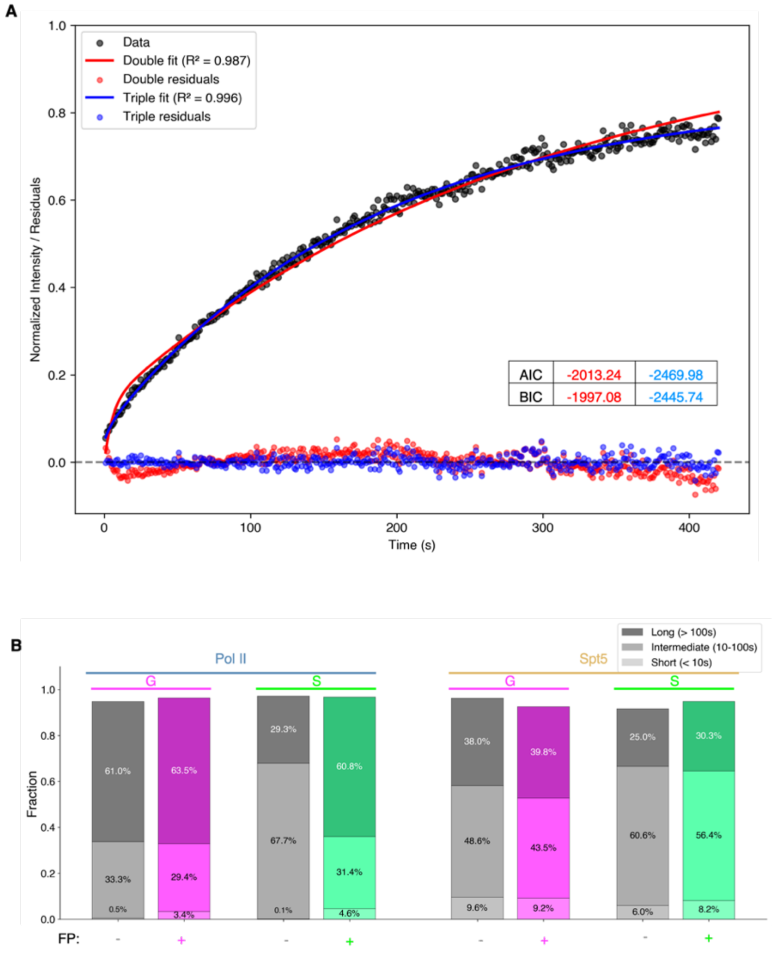

**Fig. S6. Fitting of FRAP recovery curves at HLBs reveals three kinetic components for Pol II and Spt5.** Related to **Fig. 3**. (**A**) Representative fit of a FRAP recovery curve (Pol II G-phase at HLBs; data: black; n = 13). Recovery data were fitted with sums of exponentials (red line, sum of 2 exponentials; blue line: sum of 3 exponentials; see Methods) to determine the minimal model complexity required to fit the data. Residuals are represented in the lower plot (red, double-exponential; blue, triple-exponential). The double-exponential fit produced temporally correlated residuals, whereas the triple-exponential fit eliminated systematic deviations, suggesting that three exponentials are required to describe the data. Consistent with this observation, Akaike (AIC) *(92)* and Bayes–Schwarz/Bayesian (BIC) *(93)* information criteria (inset table), which penalize additional model parameters, were lower for the triple-exponential model than for the double-exponential model, demonstrating optimal fitting by the triple-exponential model, consistent with previous work *(98)*. (**B**) Constrained multi-dataset fitting at HLBs using the same framework as in (A), with the additional constraint that one kinetic component is shared between Pol II and Spt5 (see Methods). Three kinetic components (short, intermediate, and long) were identified, and their fractional contributions are shown. In G phase, Pol II exhibits a higher fraction of the long-lived component and a lower fraction of the intermediate component than in S phase; FP shifts the S-phase species distribution toward a G-phase–like composition.

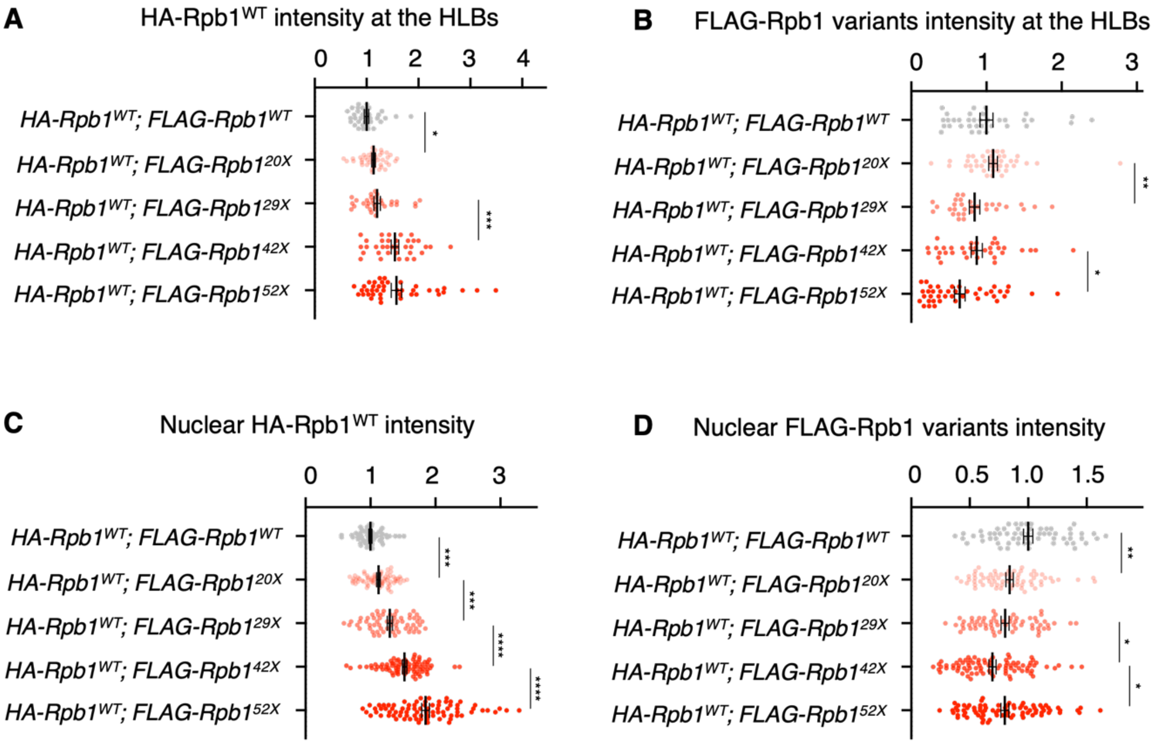

**Fig. S7. Clustered Pol II levels at HLBs do not increase at longer CTD lengths.** Related to **Fig. 4A**. (**A** and **B**) Quantifications of the raw intensity of HA-Rpb1$^{WT}$ (A) and FLAG-Rpb1 (B) at HLBs. The baseline is set to 1 for the WT. Each dot represents a nucleus. (**C** and **D**) Quantifications of the nuclear intensity of HA-Rpb1$^{WT}$ (C) and FLAG-Rpb1 (D). The moderate increase in the raw levels of HA-Rpb1WT at HLBs observed in (A) is explained by a matching increase in its nuclear levels shown in (C). Each dot represents a nucleus. Data was acquired from 4 stage-matched egg chambers from 4 individual females per genotype. All error bars represent mean ± SEM. Statistical comparisons were performed on neighboring genotypes. *$P$<0.05, **$P$<0.01, ***$P$<0.001, and ****$P$<0.0001.

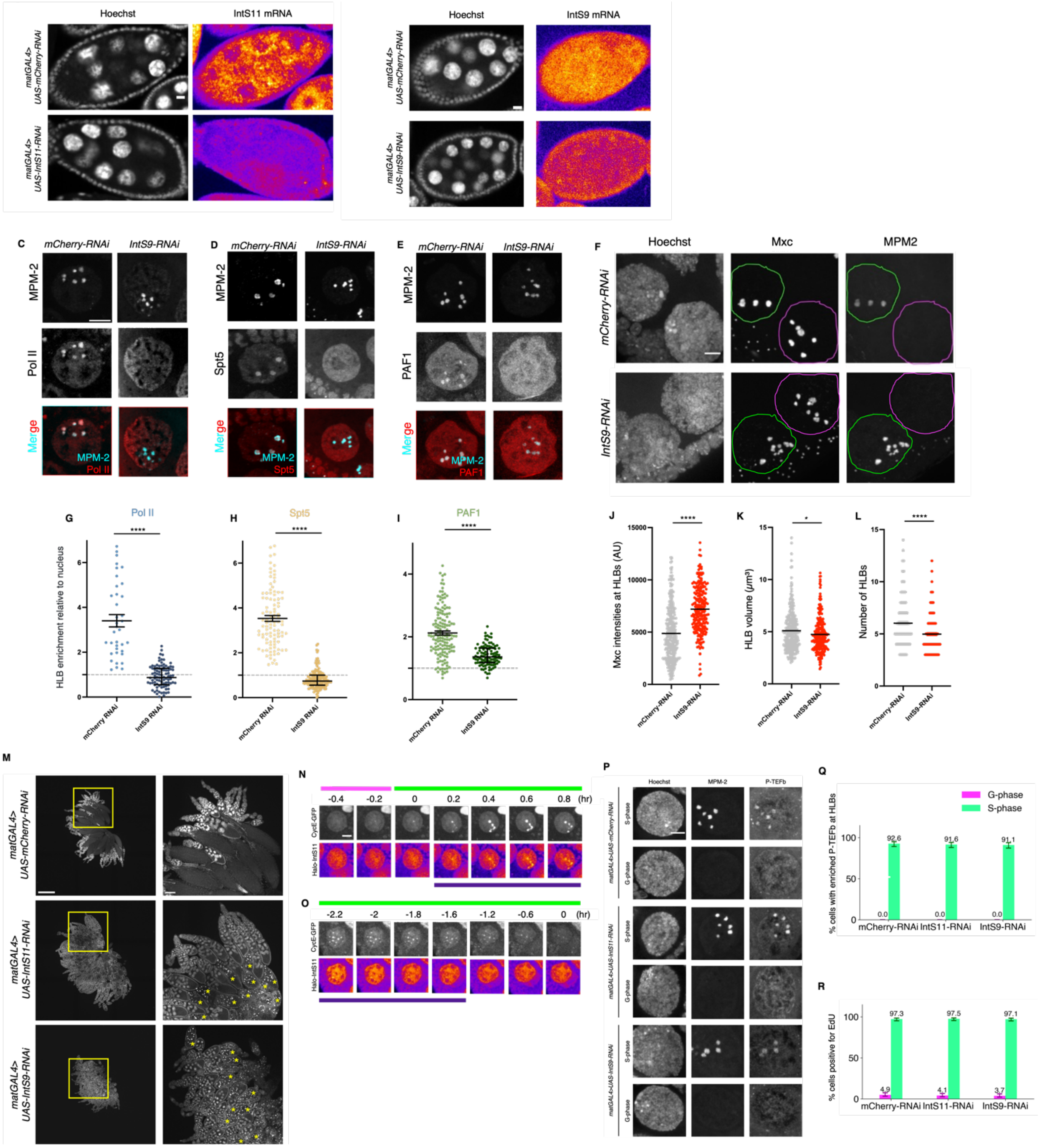

**Fig. S8. Integrator endonuclease depletion disrupts clustering without abolishing S-phase-specific P-TEFb enrichment.** Related to **Fig. 5**. (**A** and **B**) Validation of IntS11 (A) and IntS9 (B) RNAi efficiencies using RNA FISH. shRNA expression was driven by mat-GAL4. mCherry-RNAi serves as a negative control. Scale bars, 10 µm. (**C** to **F**) RNAi depletion of IntS9 results in a loss of Pol II (C), Spt5 (D), and PAF1 clusters (E) at HLBs, whereas Mxc levels are not reduced (F). mCherry-RNAi serves as a negative control. Scale bars, 10 µm. (**G** to **L**) Quantification of Pol II (G), Spt5 (H), PAF1 (I), and Mxc (J to L) at HLBs. Pol II, Spt5, and

PAF1 (G to I) intensities at HLBs (demarcated by MPM-2) were normalized to nucleoplasm levels. For Mxc (J to L), HLB intensity, HLB volume, and number of HLBs per NC were measured based on Mxc staining (F). Each dot represents one NC. Error bars, mean ± SEM. All data were stage-matched and acquired from at least 6 flies per condition. ****$P < 0.0001$. (**M**) Hoechst-stained adult *Drosophila* ovaries. Maximum Intensity Projections. The right panels are magnified versions of the boxed regions in the left panels. Yellow asterisks indicate degenerating egg chambers with distorted nuclei. Developmental arrest was observed as early as stage 10 for IntS11 RNAi and stage 9 for IntS9 RNAi; therefore, all analyses were restricted to stages 6–9 for IntS11 RNAi and stages 6–8 for IntS9 RNAi. Scale bars, 500 µm (left) and 100 µm (right). (**N** and **O**) Representative time-lapse snapshots of Halo-IntS11 in nuclei entering (N) or exiting (O) S phase. S-phase nuclei are identified by enrichment of CycE–GFP at the HLBs. Time 0 denotes the onset of S phase (entering panels) or the onset of G phase (exiting panels). Halo-IntS11 was labeled with Janelia Fluor 549. Top magenta bar marks G phase frames, top light green bar marks S phase frames; bottom color bar marks frames where the factor is detected at the HLB. Scale bar, 10 µm. (**P**) P-TEFb remains enriched at HLBs in an S-phase-specific manner upon RNAi depletion of IntS11 and IntS9. MPM-2 serves as a marker of S-phase. DNA was detected by Hoechst as a reference for the nuclei. Scale bar, 5 µm. (**Q**) Percentage of NCs positive for P-TEFb at HLBs (S-phase vs. G-phase, as determined by MPM-2 staining). $n$ >250 NCs per cell-cycle phase. **(R)** Percentage of NCs positive for EdU (S-phase vs. G-phase, as determined by MPM-2 staining) to validate the accuracy of cell-cycle staging upon RNAi depletion of IntS11 and IntS9. $n$>400 NCs per cell-cycle phase.

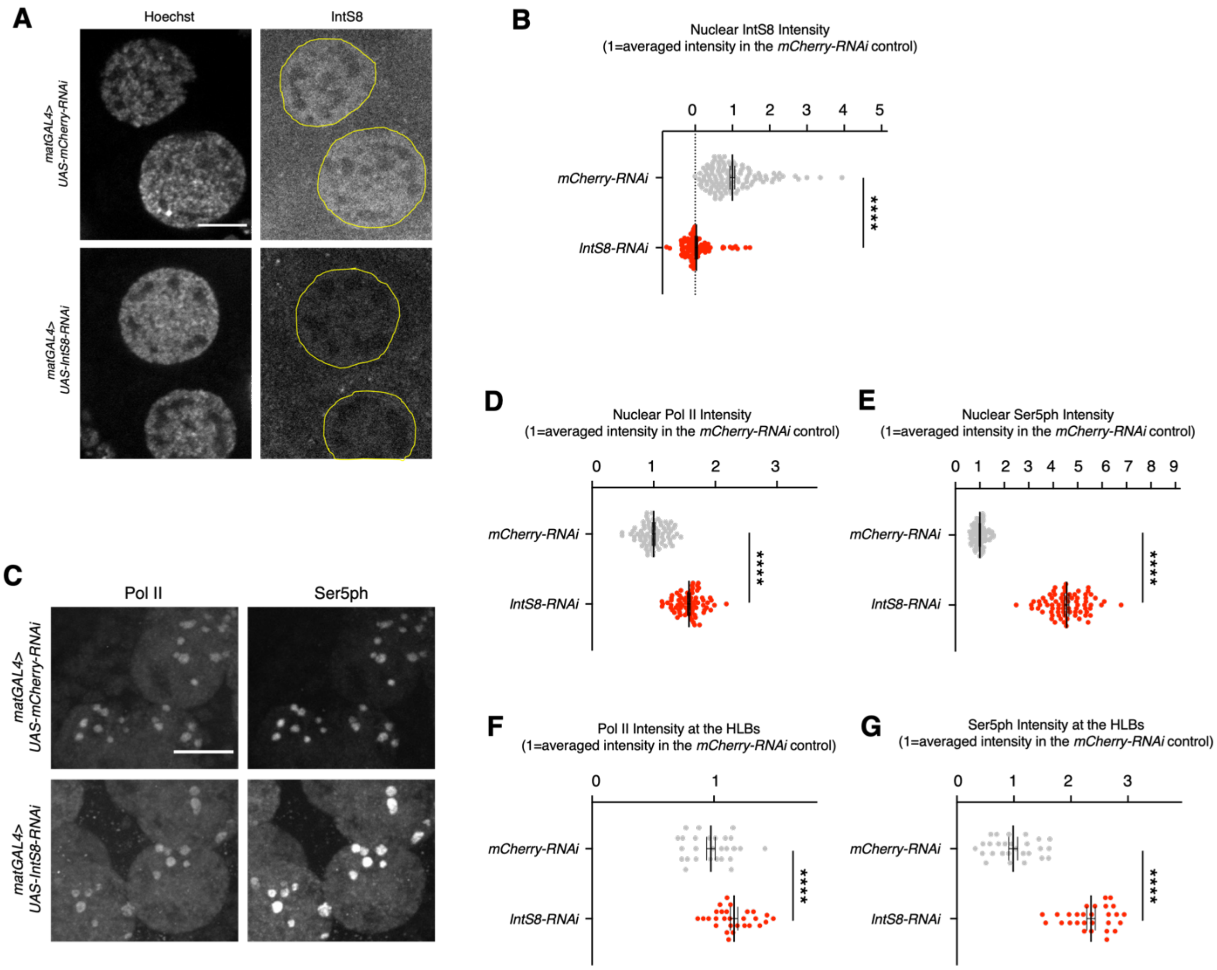

**Fig. S9. RNAi depletion of IntS8 leads to increased Ser5ph without reducing the levels of Pol II at HLBs.** (**A**) IntS8 RNAi efficiently depletes nuclear IntS8. Pol II and IntS8 were detected using antibodies against Rpb1 (ARNA3) and the *Drosophila* IntS8, respectively (85). Nuclei are outlined by the yellow circles. (**B**) Quantifications of nuclear IntS8 intensities. Each dot represents a nucleus. Error bars represent mean ± SEM. Data was acquired from 8 stage-matched egg chambers (3 females) per genotype. (**C**) IntS8 RNAi leads to drastically increased Ser5ph levels both in the overall nuclei and at HLBs without dissolving Pol II clusters. (**D** to **G**) Quantifications of Pol II and Ser5ph intensities in the nuclei (D and E) and at HLBs (F and G). The increased levels of Ser5ph are not explained by levels of Pol II. Nuclear: 4.56±0.09 (Ser5ph) versus 1.57±0.02 (Pol II); HLBs: 2.38±0.08 (Ser5ph) versus 1.20±0.03 (Pol II). Each dot represents a nucleus. Error bars represent mean ± SEM. Data were acquired from 5 stage-matched egg chambers (3 females) per genotype. Scale bars, 10 µm. ****$P<0.0001$.

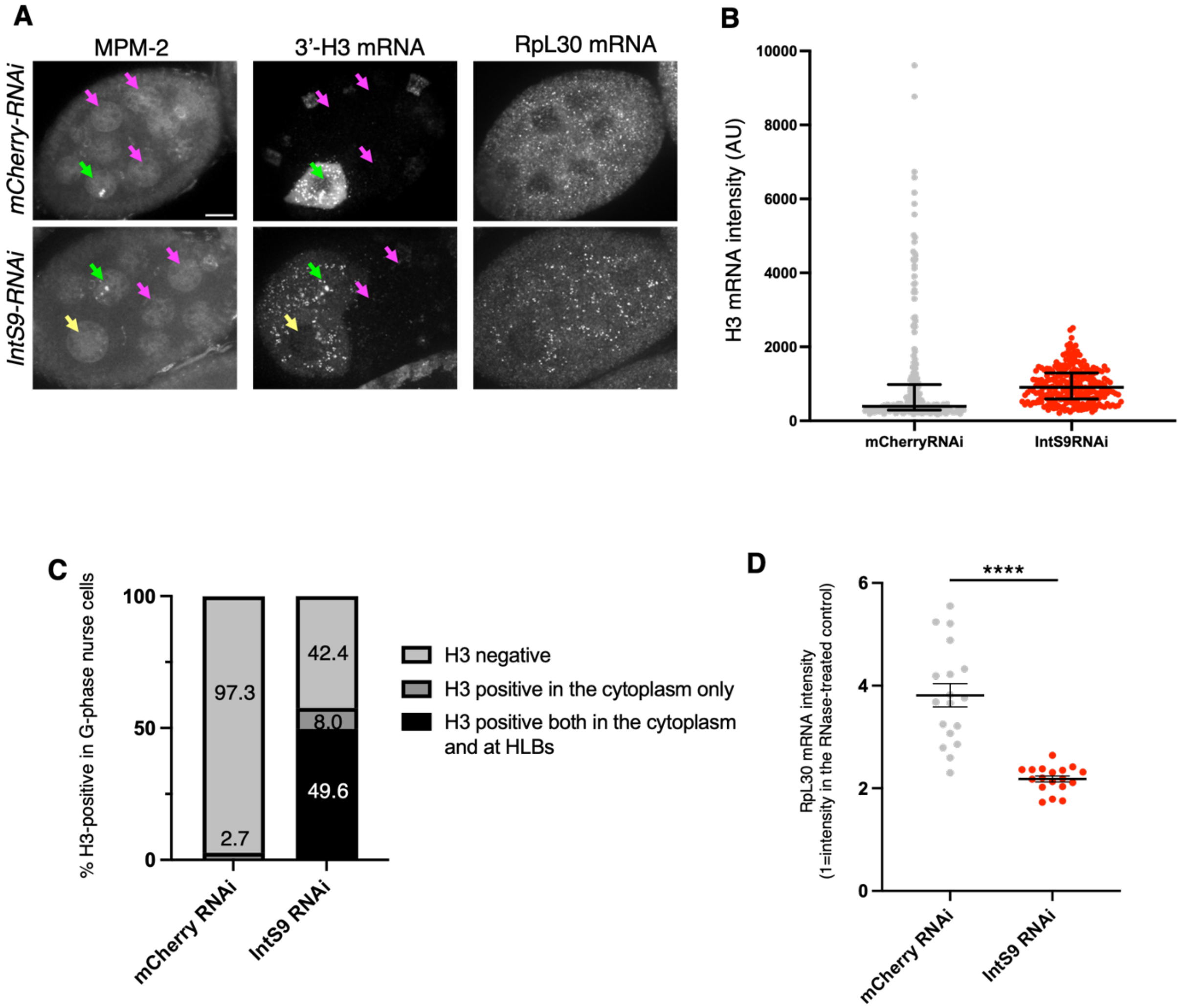

**Fig. S10. Depletion of IntS9 results in reduced induction and out-of-phase histone transcription.** Related to **Fig. 6**. (**A**) Egg chambers probed with MPM-2 (S-phase marker), 3'-H3 mRNA, and RpL30 mRNA (control). mCherry-RNAi serves as a negative control. Green arrows indicate S-phase NCs, whereas magenta and yellow arrows indicate G-phase NCs lacking and containing H3 mRNA, respectively. Scale bar, 10 µm. (**B**) Cytoplasmic H3 mRNA intensity. Each dot represents one NC. Bars indicate quartiles. $P<0.00001$, estimated by bootstrapping of the 10$^{th}$ and 90$^{th}$ percentiles. (**C**) Percentages of MPM-2-negative NCs with no H3 mRNA signal, H3 mRNA in the cytoplasm only, and H3 mRNA both in the cytoplasm and at HLBs. $n > 300$ NCs for each genotype. (**D**) Cytoplasmic RpL30 mRNA FISH intensity. Each dot represents the average mRNA level in one egg chamber. Error bars, mean ± SEM.

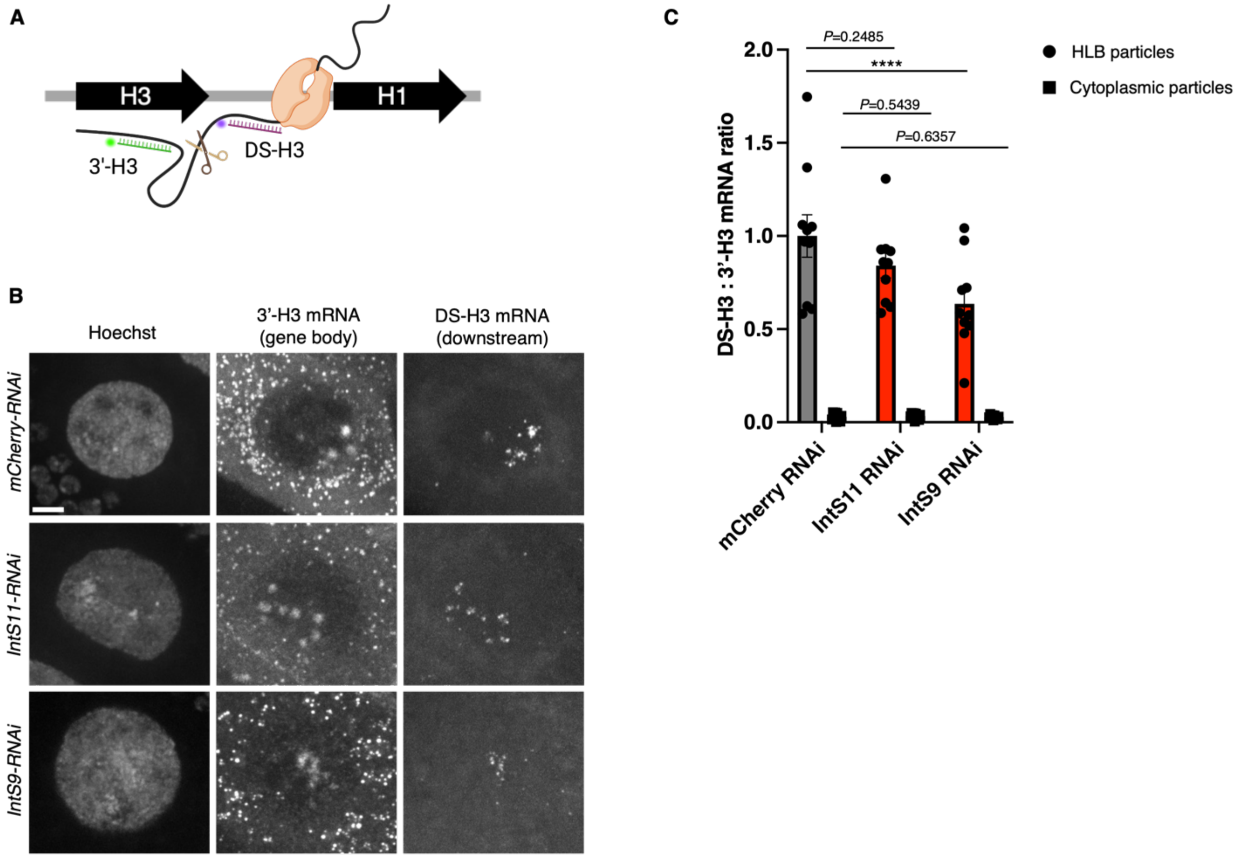

**Fig. S11. Integrator endonuclease depletion does not impair histone H3 mRNA processing.** Related to **Fig. 6**. (**A**) Schematic of the downstream H3 (DS-H3) probe for the detection of unprocessed H3 mRNA, adapted from(*42*). Scissors indicate the mRNA cleavage site. (**B**) Representative images showing NCs probed with both the 3'-H3 and DS-H3 FISH probes. DNA was detected by Hoechst as a reference for the nuclei. Data were stage-matched and collected from at least six females per genotype. Scale bar, 10 µm. (**C**) Quantification of DS-H3 to 3'-H3 mRNA ratios at HLBs and the mRNA clusters in the cytoplasm. Each dot represents one NC. Error bars represent mean ± SEM.

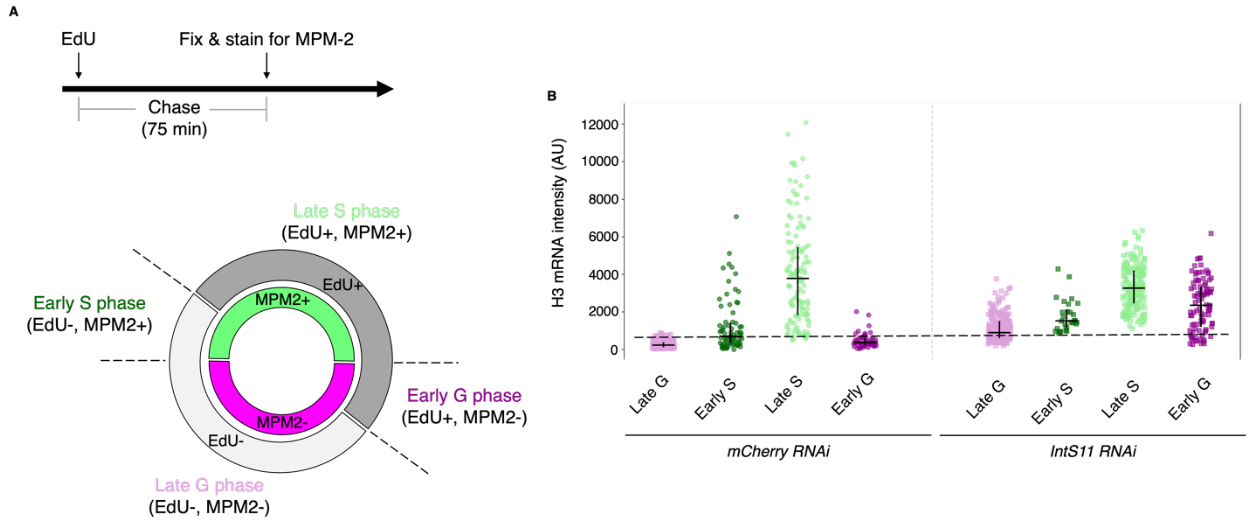

**Fig. S12. Loss of IntS11 triggers premature activation of histone transcription.** Related to **Fig. 6**. (**A**) Schematic of EdU/MPM-2 combinatorial labeling for refined cell-cycle staging. To better resolve transitions between cell-cycle phases, egg chambers were pulsed with EdU for 15 min, chased in EdU-free medium for 75 min, and subsequently labeled for MPM-2. This strategy allows the NC cell cycle to be partitioned into four distinct stages based on EdU and MPM-2 status. (**B**) Quantification of *H3* mRNA levels across cell-cycle subphases. Each dot represents one NC. Bars indicate quartiles. In *mCherry-RNAi* controls, a fraction of early S-phase NCs lack detectable H3 signal, consistent with a short delay between S-phase entry and the onset of histone transcription. In contrast, *IntS11-RNAi* NCs show detectable mRNA levels across the entire early S-phase population, indicating that productive histone transcription is initiated prior to the G/S transition.

| Probe name | Probe sequence |
|---|---|
| 5'-H3-i2-F | CCTCgTAAATCCTCATCAAAACTACTTCACGTTTGAAAACACAAT |
| 5'-H3-i2-R | ATTTGGGTTTCACTAAAGTTCACGTAAATCATCCAgTAAACCgCC |
| 3'-H3-i1-1F | gAggAgggCAgCAAACggAACGAAGAGACCAACCAGGTAGGCTTC |
| 3'-H3-i1-1R | CATGAATGGCACACAAGTTGGTATCTAgAAgAgTCTTCCTTTACg |
| 3'-H3-i1-2F | gAggAgggCAgCAAACggAAAACTGGATGTCTTTGGGCATTATGG |
| 3'-H3-i1-2R | GCACGCTCGCCGCGAATGCGTCGCGTAgAAgAgTCTTCCTTTACg |
| 3'-H3-i1-3F | gAggAgggCAgCAAACggAAGCTTTATCTGCAAGTTAATGCCGTG |
| 3'-H3-i1-3R | AAAGGACCGATTATAGAGTACGCTATAgAAgAgTCTTCCTTTACg |
| RpL30-i2-A-F | CCTCgTAAATCCTCATCAAAGCCCAATTGTAATGGAAAGGGAATT |
| RpL30-i2-A-R | TGTGTTTGTCTACGTAATGTATATTAAATCATCCAgTAAACCgCC |
| RpL30-i2-B-F | CCTCgTAAATCCTCATCAAATGATGATAATCCCAACATTGCTTTC |
| RpL30-i2-B-R | ACAACAAGAATCACGTGCCAAATCTAAATCATCCAgTAAACCgCC |

| | |
|---|---|
| RpL30-i2-C-F | CCTCgTAAATCCTCATCAAATGTTTAGGCCGTCTCCAGCGAGCGG |
| RpL30-i2-C-R | GTGTCGTCTTGTTCGATTAAAAATAAAATCATCCAgTAAACCgCC |
| RpL30-i2-D-F | CCTCgTAAATCCTCATCAAAGCACACGCGGAAGTATTTACCACAG |
| RpL30-i2-D-R | ATCTCCAGGATCGGTGATGGACAGGAAATCATCCAgTAAACCgCC |
| RpL30-i2-E-F | CCTCgTAAATCCTCATCAAAAGTATTTGCCGGACTTCATCACCAG |
| RpL30-i2-E-R | TCTTCAAGGTCTGCTTGTAGCCCAGAAATCATCCAgTAAACCgCC |
| RpL30-i2-F-F | CCTCgTAAATCCTCATCAAAACTTCGTCGGCTGACAATGGCAAAA |
| RpL30-i2-F-R | CACCATGTTGATAGTTTGGGTTAAAAAATCATCCAgTAAACCgCC |
| IntS11-i3-1F | gTCCCTgCCTCTATATCTTTTTCTGGAAGGGATCAGGATCACCCG |
| IntS11-i3-1R | CAAGGGCGTTATCTTGATGTCCGGCTTCCACTCAACTTTAACCCg |
| IntS11-i3-2F | gTCCCTgCCTCTATATCTTTGTGTCCACCATCATACTCTGGTGAA |
| IntS11-i3-2R | GCATAGTAGGCTTTGATCTCAAGGTTTCCACTCAACTTTAACCCg |
| IntS11-i3-3F | gTCCCTgCCTCTATATCTTTGTATATGGGGTACTTGAGGTTCATG |
| IntS11-i3-3R | GGCCTTCTCGGTAAGACCCAGAGCATTCCACTCAACTTTAACCCg |
| IntS11-i3-4F | gTCCCTgCCTCTATATCTTTCTTTGGCTCACAGTTCTGAATTAGC |
| IntS11-i3-4R | TGCTTCACCATGGACGAGCATGACGTTCCACTCAACTTTAACCCg |
| IntS11-i3-5F | gTCCCTgCCTCTATATCTTTATTCGATTGTCCTTCATTACGAGGA |
| IntS11-i3-5R | AGGGCATCGGTTAGGTTTTGCAGCATTCCACTCAACTTTAACCCg |
| IntS11-i3-6F | gTCCCTgCCTCTATATCTTTCACATTCAGTATGTAGGCTCCAATG |
| IntS11-i3-6R | CTTCGTTGCCTAGCACATATTCTGCTTCCACTCAACTTTAACCCg |
| IntS9-i3-A-F | gTCCCTgCCTCTATATCTTTCATTGACAAGGTTCAGAGTCAGAGC |
| IntS9-i3-A-R | TGTCCTCGAACTTAATGACTGTGTCTTCCACTCAACTTTAACCCg |
| IntS9-i3-B-F | gTCCCTgCCTCTATATCTTTCAGTGGACCTTGTCCTTAACTTGCA |
| IntS9-i3-B-R | TTAACGCTATCCGCACATGGCTGGATTCCACTCAACTTTAACCCg |
| IntS9-i3-C-F | gTCCCTgCCTCTATATCTTTGGATGACAAGCACATTTGGTTTCAA |
| IntS9-i3-C-R | AGGGATGTGGTTTTGTGTATGCCTCTTCCACTCAACTTTAACCCg |
| IntS9-i3-D-F | gTCCCTgCCTCTATATCTTTTCTCGATAAAGTGAACAGCGTCGCC |
| IntS9-i3-D-R | TGGAGTTGTTAGGGTTATTGCCCCATTCCACTCAACTTTAACCCg |
| IntS9-i3-E-F | gTCCCTgCCTCTATATCTTTCACTCGGCCAGGATATTAGAATACG |
| IntS9-i3-E-R | ACTTTGTTCTGCTTGGCGGAACTAATTCCACTCAACTTTAACCCg |
| IntS9-i3-F-F | gTCCCTgCCTCTATATCTTTATAGACAACGCCCGAAGGATAACAG |
| IntS9-i3-F-R | ATTCTGGGTGAGGCACTCAAAGAGGTTCCACTCAACTTTAACCCg |

**Table S1. Sequences of HCR FISH probe pairs.**

| |
|---|
| CCTCgTAAATCCTCATCAAAAATTAGTATTTTTACATATGATTTA |
| AGATTATTTTTATTCCTTCTCTGGCAAATCATCCAgTAAACCgCC |

| |
|---|
| CCTCgTAAATCCTCATCAAATAATAAATGTCGAGCTTACAATTAA |
| TTATATTTATTAACCTCTTCTTTAAAAATCATCCAgTAAACCgCC |
| CCTCgTAAATCCTCATCAAATTCGCACAAAGAACAAATAAAAATA |
| ACGACACTTTCACTGCTTTAAAGGAAAATCATCCAgTAAACCgCC |
| CCTCgTAAATCCTCATCAAATGTTTAAGGTTCAGAGTCCCTTTCC |
| ACACAGAATTCAGATTTTTTTTTAAAATCATCCAgTAAACCgCC |
| CCTCgTAAATCCTCATCAAAATGTAATTTCATTAATTTCAAACTG |
| GTTATTTTATAAATATTGCCGTCATAAATCATCCAgTAAACCgCC |
| CCTCgTAAATCCTCATCAAAAAATAGCTAGTTTTATTTTATTATT |
| CTTCAGTTAACACATGGAAAAAATAAAATCATCCAgTAAACCgCC |
| CCTCgTAAATCCTCATCAAACCCGTACGACCTCTTCAATAATAAC |
| GGCGTTTGAAGGGACAGTGTCAATTAAATCATCCAgTAAACCgCC |
| CCTCgTAAATCCTCATCAAATGAATTTTACATAGGTTTTATTTTT |
| CGCAACAAAATTAGCCAATTTCCGTAAATCATCCAgTAAACCgCC |
| CCTCgTAAATCCTCATCAAATCCTTTATTATGTAATATATATTAC |
| AAATAAAAAGAAACAATTTTTGTATAAATCATCCAgTAAACCgCC |
| CCTCgTAAATCCTCATCAAATATGTAGTCAAATAAATAAATCAAA |
| TTTCCTCGCCACATATGCATTACCGAAATCATCCAgTAAACCgCC |
| CCTCgTAAATCCTCATCAAATTAAAATAATTTGTTCTGAAATCAA |
| TGTTATTAATGTGTTTTCATGCATAAAATCATCCAgTAAACCgCC |
| CCTCgTAAATCCTCATCAAAAATAACATAATATTAATATAATAAA |
| ATCGTTAAAAATACAGATACTTTCTAAATCATCCAgTAAACCgCC |
| CCTCgTAAATCCTCATCAAATTTAAAGCAGCATTCAGAAATAATT |
| TTCACATTGAGCTACAGAAAAATTTAAATCATCCAgTAAACCgCC |
| CCTCgTAAATCCTCATCAAAAATTAAGATACTTTATTACTTGATT |
| AGGCTTCTGAAAGAAGCGTCTATTAAAATCATCCAgTAAACCgCC |

| |
|---|
| CCTCgTAAATCCTCATCAAATTATTAAAATTGAAACGTTCATCCC |
| TGCAAATATTTCACTAATTCGTAATAAATCATCCAgTAAACCgCC |
| CCTCgTAAATCCTCATCAAAAAAAAAACATCTAATAAAATAAGAA |
| TTCAAATTTTTAACTGGACCAATTTAAATCATCCAgTAAACCgCC |
| CCTCgTAAATCCTCATCAAAGACTTTCAAATGTTGATTGAATTTT |
| TTTAAAAGGATGTAATATGGGTTTTAAATCATCCAgTAAACCgCC |
| CCTCgTAAATCCTCATCAAATTTAAATAATTTTTTCGTCACTTTC |
| AAAATAAGGTTTTAATAGTTCTACAAAATCATCCAgTAAACCgCC |
| CCTCgTAAATCCTCATCAAATAATTTTTTAGTATTTTAAATCATT |
| AAGTTTGCTTGAAGTGTAAACTTTTAAATCATCCAgTAAACCgCC |
| CCTCgTAAATCCTCATCAAATGCTACTGACATCAGTCATTTACTA |
| GAATCGAGGAGGAGAGCACTTTAACAAATCATCCAgTAAACCgCC |

**Table S2. Sequences of HCR FISH probe for DS-H3.**

**Movie S1. Pol II clusters form at HLBs during the G-phase.** Live movie of an egg chamber expressing CycE-GFP and Halo-Rpb1, the former localizes to HLBs during the S-phase. Halo-Rpb1 was labeled by JF549. The boxed region corresponds to snapshots shown in **Fig. 1G**, which captures an NC entering the S-phase. Time 0 represents the transition into the S-phase. Scale bars, 10 μm.

**Movie S2. Pol II clusters persist at HLBs after NCs exit the S-phase.** Live movie of another egg chamber expressing CycE-GFP and Halo-Rpb1. Halo-Rpb1 was labeled by JF549. The boxed region corresponds to snapshots shown in **Fig. 1H**, which captures an NC exiting the S-phase. Time 0 represents the transition into the G-phase. Scale bars, 10 μm.

87. Babraham Bioinformatics - FastQC A Quality Control tool for High Throughput Sequence Data. https://www.bioinformatics.babraham.ac.uk/projects/fastqc/.

88. *Trim Galore: A Wrapper Tool around Cutadapt and FastQC to Consistently Apply Quality and Adapter Trimming to FastQ Files*.
